# Neuron Enriched Exosomal MicroRNA Expression Profiles as a Marker of Early Life Alcohol Consumption

**DOI:** 10.1101/2023.06.09.544235

**Authors:** Vasily Yakovlev, Dana M. Lapato, Pratip Rana, Preetam Ghosh, Rebekah Frye, Roxann Roberson-Nay

## Abstract

**Background:** Alcohol consumption may impact and shape brain development through perturbed biological pathways and impaired molecular functions. We investigated the relationship between alcohol consumption rates and neuron-enriched exosomal microRNA (miRNA) expression to better understand the impact of alcohol use on early life brain biology.

**Methods:** Neuron-enriched exosomal miRNA expression was measured from plasma samples collected from young people using a commercially available microarray platform while alcohol consumption was measured using the Alcohol Use Disorders Identification Test. Linear regression and network analyses were used to identify significantly differentially expressed miRNAs and to characterize the implicated biological pathways, respectively.

**Results:** Compared to alcohol naïve controls, young people reporting high alcohol consumption exhibited significantly higher expression of four neuron-enriched exosomal miRNAs including miR-30a-5p, miR-194-5p, and miR-339-3p, although only miR-30a-5p and miR-194-5p survived multiple test correction. The miRNA-miRNA interaction network inferred by a network inference algorithm did not detect any differentially expressed miRNAs with a high cutoff on edge scores. However, when the cutoff of the algorithm was reduced, five miRNAs were identified as interacting with miR-194-5p and miR-30a-5p. These seven miRNAs were associated with 25 biological functions; miR-194-5p was the most highly connected node and was highly correlated with the other miRNAs in this cluster.

**Conclusions:** Our observed association between neuron-enriched exosomal miRNAs and alcohol consumption concurs with results from experimental animal models of alcohol use and suggests that high rates of alcohol consumption during the adolescent/young adult years may impact brain functioning and development by modulating miRNA expression.

## Introduction

Adolescents and young adults frequently consume alcohol and often engage in binge drinking, making it the most used and misused substance among teens^1–3^. Although most young people who binge or heavily drink do not currently have an alcohol use disorder (AUD), these alcohol use (AU) patterns can lead to lasting neurobiological changes and an increased likelihood of developing an AUD^4–11^. The cellular processes underlying the effects of alcohol use and misuse in young people are not well understood.

A diverse array of molecular modifications, nucleic acids, and cellular machinery jointly regulate gene expression. Much attention has been paid to characterizing the intracellular biological pathways and actors that augment or repress messenger RNA (mRNA) abundance and protein production, which has led to an increased appreciation of the role of microRNAs. MicroRNAs (miRNA) are non-coding RNAs that coordinate with cellular machinery to finetune gene expression^12^. MiRNAs exhibit self-regulatory mechanisms, regulate the expression of other genes/miRNAs, and influence biological processes within cells. Although miRNA activity is present throughout the entire human life course, miRNAs play especially key roles during development, differentiation, and cell-fate determination^12^. Additional evidence suggests that miRNAs serve as key roles in immune system functioning, aging, and reproduction.

Several preclinical studies and a few human studies have investigated microRNAs (miRNAs) in the context of alcohol use. These studies reveal miRNAs that map to genes related to hypothesized etiologic alcohol use pathways, including neuroplasticity/brain development and immune system/inflammation^13–23^. For instance, in human post-mortem brain samples, approximately 35 miRNAs exhibited differential expression levels in the prefrontal cortex (PFC) of adult humans who abused alcohol compared to controls and over-expression of miRNAs was inversely associated with the expression of their target mRNAs^24^. Robust changes in miRNA expression also have been reported in the brains of rodents following chronic alcohol exposure, with ∼41 rat miRNAs being significantly altered in the medial PFC^25^. Gene ontology results indicated that differential expression was related to functional processes commonly associated with neurotransmission, synaptic plasticity, and neuroadaptation. In a study comparing persons with AUDs to non-drinking controls, increased expression was found for mir-96, mir-24, and mir-136 and decreased mir-92b expression in persons with AUDs^26^. These findings are consistent with human and animal studies of brain tissue^27–30^. Moreover, the observation of primarily increased levels among persons with AUDs is in agreement with the findings from post-mortem brain tissue derived from persons suffering from alcohol use disorder^24^ and suggests the presence of a potential compensatory mechanism in response to neuronal damage or, possibly, an intensified transfer of miRNAs from the brain to the peripheral regions due to cellular harm^26^. Thus, miRNA expression may hold great potential to elucidate pathways to alcohol-induced brain alterations and damage^31, 32^.

Given that all miRNAs are not thought to participate equally in every biological function or developmental process, it is not surprising that miRNAs vary considerably in expression levels^33^ and that these levels almost certainly vary by sex, age, health status, and environmental (e.g., exposure to pollution) and behavioral (e.g., substance use) factors. Most importantly, miRNAs can be shuttled through the bloodstream by extracellular vesicles (EVs). EVs are robust biological sacs that transport cargoes of proteins, RNAs (including miRNAs), DNA, and lipids throughout the body and across the blood-brain barrier. This portability enables brain-derived EVs to be isolated from peripheral blood, opening a window to the brain by identifying, quantifying, and characterizing EV cargo throughout the lifespan to understand both normal physiology and pathophysiology^34^.

The current study investigated the abundance and potential roles of circulating exosomes during adolescence/young adulthood. Exosomes are a subclass of EVs that have been associated with a wide range of conditions, including neurodegenerative disorders and cancer ^35–40^. The majority of EV and exosome research has focused exclusively on adult populations, leaving significant knowledge gaps regarding the number, type, and cargo content of exosome in healthy children, adolescents, and young adults and whether perturbations in any of these characteristics could be concomitant with, predict, or follow significant changes in health, development, or functioning. For the cargo analysis, we prioritized miRNAs in the EV cargo analysis given that miRNAs serve prominent roles in brain functioning and development^41^, are relatively abundant in many EVs, and are amenable to high throughput measurement. Additionally, miRNAs are inherently biologically active. Unlike mechanisms like DNA methylation, which may or may not be directly impacting expression profiles of the genic regions it overlaps, miRNAs are mobile and capable of associating with targets and partner cellular machinery from the moment that they exit EVs and enter recipient cells. Moreover, robust methods and resources^33^ exist to identify, quantify, and characterize miRNAs and their functionality, making miRNAs more tractable candidates for gene network analyses than more enigmatic and less well-characterized options like long noncoding RNAs.

## Methods

### Participants

All specimens and participant data used in this study were collected as part of the Adolescent and Young Adult Twin Study (AYATS; R01MH101518; IRB protocol #HM20022942) and recruited primarily through the Mid-Atlantic Twin Registry. AYATS included two study visits separated by two years. Young people reporting high alcohol consumption rates (i.e., AUDIT-C ≥ 6) were selected for inclusion. Given that alcohol use rates are highly correlated among twin pairs, this resulted in 5 twin pairs being included in the AU case group, with the remaining 12 alcohol cases being individuals. All control participants were individuals (i.e., no twin pairs).

During each visit, participants completed self-report questionnaires and provided blood samples. Self-report questionnaires queried youth regarding multiple domains including psychiatric conditions, personality, social relations, and substance use. Blood samples were spun down to separate the plasma, which was stored at −80℃.

### Alcohol Consumption

The Alcohol Use Disorders Identification Test (AUDIT) was developed by the World Health Organization and is the most widely used alcohol screening instrument of past-year alcohol use patterns^42–50^. The test consists of 10 items across three dimensions including consumption (termed AUDIT-C), dependence, and problematic or hazardous alcohol use. A meta-analysis found the 3-item AUDIT-C to be as effective as the full 10-item AUDIT in screening for at-risk drinking and AUDs. Responses to each of the three questions are assigned 0-4 points, yielding a total score of 0-12. AUDIT-C screening thresholds that optimize sensitivity and specificity for detecting unhealthy alcohol use are ≥4 in males and ≥3 in females^44^ whereas a study of adolescents suggested a cutoff score of ≥3 as indicative of problematic alcohol use for males and females^50^. AUDIT-C scores will serve as the primary AU variable given that we are most interested in the impact of alcohol consumption rates on brain neurobiology of young people.

### Short Mood and Feelings Questionnaire (SMFQ)

The SMFQ is a 13-item questionnaire that maps onto DSM criteria for depression, with a total range of 0-26. The time frame for answering the SMFQ is the *<u>past two weeks</u>* and a total score greater than 12 may indicate the presence of depression. The SMFQ is a well validated measure normed for adolescents^51–53^.

### Smoking Behaviors

Use of combustible cigarettes, electronic cigarettes, and marijuana was assessed. Given that no one smoking behavior was endorsed with high frequency, all three behaviors were aggregated to generate a single index of smoking behaviors. The smoking behavior variable was coded as binary (0=no smoking behaviors endorsed, 1=one or more smoking behaviors endorsed).

### Exosome Isolation and Enrichment

Exosomes were extracted from plasma samples by size exclusion chromatography method using the qEVoriginal columns (Izon Science). Every 0.5 ml of plasma yielded 1.5 ml of purified exosomes in the 1xPBS buffer. Exosome precipitation was performed overnight at +4℃ on a rotating mixer with L1CAM/CD171 Monoclonal Biotin Conjugated Antibody (eBio5G3 (5G3), Thermofisher) in the presence of 3% BSA and 3x Proteinase/Phosphatase Inhibitors cocktail (Sigma-Aldrich). After overnight incubation samples were additionally incubated with Pierce™ Streptavidin Plus UltraLink™ Resin (Thermofisher) for 1 hour at +4℃. As a washing step, sample tubes were centrifuged with 500xg for 1 min, the supernatant was removed and saved as non-neuronal fraction of plasma exosomes, and resin beads were washed 3x times with cold 1 ml of pre-filtered (with 0.22μM filter) 1xPBS buffer. After final wash, precipitated neuronal-enriched exosomes were eluted by incubation of the magnetic beads with 100 μl of Pierce™ IgG Elution Buffer (ThermoFisher) at room temperature with mixing for 5 minutes.

### Nanoparticle tracking analysis

The concentration and size of exosomes were measured using ZetaView® Nanoparticle Tracking Analysis. All samples were diluted in PBS to a final volume of 2 ml. Ideal measurement concentrations were found by pre-testing the ideal particle per frame value (140–200 particles/frame). The manufacturer’s default software settings for exosomes were selected accordingly. For each measurement, three cycles were performed by scanning 11 cell positions each and capturing 80 frames per position under following settings: Focus: autofocus; Camera sensitivity for all samples: 78; Shutter: 100; Scattering Intensity: detected automatically; Cell temperature: 25°C. After capture, the videos were analyzed by the in-built ZetaView Software 8.04.02 SP2 with specific analysis parameters: Maximum area: 1000, Minimum area 5, Minimum brightness: 25. Hardware: embedded laser: 40 mW at 488 nm; camera: CMOS. The number of completed tracks in NTA measurements was always greater than the proposed minimum of 1000 to minimize data skewing based on single large particles.

### Western blot analysis

For protein analysis all samples were loaded and separated by SDS-PAGE gel and transferred to nitrocellulose membranes. The membranes were exposed to antibodies at specific dilutions. Primary antibodies used for WB: anti-L1CAM(CD171) (dilution 1:1000, ThermoFisher), anti-CD63 (dilution 1:1000, ThermoFisher), anti-TSG101 (dilution 1:500, Cell Signaling), anti-Albumin (dilution 1:1000, Cell Signaling). Specific protein bands were detected using infrared-emitting conjugated secondary antibodies: anti-rabbit DyLight™ 800 4X PEG Conjugate (dilution 1:10,000, Cell Signaling). WB images were generated and analyzed using the ChemiDoc Infrared Imaging System (Bio-Rad).

### Transmission Electron Microscopy (TEM)

Isolated exosomes were fixed and prepared for the TEM as was previously described^54^. The TEM analysis was carried out by the Microscopy Core of VCU.

### Exosomal RNA Isolation and RT-PCR

Total RNA was isolated from the exosomal samples following the manufacturer’s instructions with the miRNeasy MicroRNA Extraction Kit (Qiagen) and then converted to cDNA using the miRCURY LNA RT Kit (Qiagen). The samples with total exosomal RNA and cDNA were stored in a freezer at −80℃. The concentration and purity of the exosomal RNA samples were estimated using NanoDrop ND-1000 spectrophotometer (Thermo Scientific, Wilmington DE).

All cDNA samples were pooled into two groups corresponding to cases (alcohol drinkers) and controls (non-drinkers) and then assayed using the Serum/Plasma Focus PCR panel (Qiagen). This commercially available microarray platform contains 179 LNA miRNA primer sets of miRNAs that have been commonly found in human plasma. The Serum/Plasma Focus miRNA PCR Panels include potential reference genes and probes for evaluating potential sample contamination or damage (e.g., hemolysis). Each PCR panel also contains set of the negative control (H_2_O) and five sets of RNA Spike-In controls to evaluate RNA extraction (Spike-Ins 2-4-5; concentration ratio 1:100:10000) and cDNA synthesis (Spike-In 6) and to perform inter-plate calibration (Spike-In 3). The amplification was performed in a QuantStudio 5 Real-Time PCR System (Thermo Fisher Scientific, USA). Samples were amplified using RT2 SYBR® Green qPCR Mastermix with ROX (carboxy-X-rhodamine) passive reference dye from QIAGEN. The real-time PCR data were normalized by ROX passive reference. The amplification curves were analyzed using the QuantStudio™ Design & Analysis Software v1.4.2, both for determination of Ct values and for melting curve analysis. All assays were inspected for distinct melting curves and the Tm was checked to be within known specifications for each assay. The 2-ΔΔCt method was used for the calculation of relative miRNA expression levels.

### RT-PCR Data Analysis

RT-PCR count data was imported into the R statistical environment^55^. Potential reference genes were identified by comparing the overall stability and relative expression of candidate miRNAs across cases and controls. MiRNAs that exhibited at least a 5-fold difference in expression between cases and controls were subsequently assayed in all individual participants using quantitative PCR. The individual-level miRNA expression values were imported in the R statistical environment where 21 multivariate linear regression analyses were conducted to test for an association between AU group and miRNA expression. Demographic factors (i.e., age, sex) served as covariates. We also included smoking behaviors and current depression symptoms as covariates given that these behaviors are frequently concomitant with AU in teens^56–66^ and we included family id as a fixed effect to account for twin pair relatedness. Partial eta squared (η^2^) was computed for all variables in the model. This effect size estimate reflects the proportion of variance explained by a variable in relation to the total variance, while accounting for the variance explained by the other variables in the model. A Bonferroni correction also was applied to correct for multiple testing (21 tests, alpha =0.05; Bonferroni adjusted *p*≤0.002).

### Functional Network Analysis

We applied a network-based approach to analyze miRNA regulation and find potential miRNA biomarkers of alcohol consumption. We used the miRsig tool^67, 68^ to infer the miRNA-miRNA interaction network. MiRsig tool operates by 1) collecting miRNAs expression in a disease/condition from multiple patients, 2) running six reverse-engineering network inference algorithms on the expression data to infer miRNA-miRNA interaction scores/probability, and 3) performing a consensus-based approach to get a more accurate network from these six miRNA-miRNA interaction algorithm outputs. Each network inference algorithm has some individual bias, and their performance varies in different datasets. The consensus approach reduces the noise and prediction error by employing a consensus of six different algorithms as demonstrated by Nalluri et al.^67^.

## Results

### Participants

Table 1 provides summary statistics for demographic, smoking, and alcohol use variables by alcohol case-control status. Of the 44 participants selected for study inclusion, n=26 (59%) endorsed alcohol consumption while n=18 self-reported as not having initiated alcohol use in their lifetime. Two individuals were excluded from the analyses, with one excluded due to poor sample quality and the second participant for a nonqualifying AUDIT-C score (i.e., AUDIT-C=2). These exclusions resulted in a final sample of 42 young people, with 24 classified as AU cases and the remainder as alcohol naïve controls. AU cases reported high rates of alcohol consumption (i.e., Mean AUDIT-C = 8.0, SD = 1.0, range 6-10). The alcohol use groups did not differ on sex, race, ethnicity, or current depression symptoms (see Table 1). Young people classified as AU cases were older (t=-3.7, p=.001) and significantly more likely to endorse smoking behaviors, with approximately 42% reporting one or more smoking behaviors (χ^2^=9.8, p=002).

**Table 1.** Demographic and clinical features of alcohol use groups.

|  | **Alcohol Use Group** |  | **t/$\chi^2$** | **p** |
| --- | --- | --- | --- | --- |
|  | **Controls** | **Cases** |  |  |
| Age | 18.1 (1.39) | 19.7 (1.5) | -3.7 | 0.001 |
| Sex, n (%) female | 9 (50.0) | 8 (33.3) | 1.2 | 0.276 |
| Race, n (%) Caucasian | 18 (100) | 24 (100) | --- | --- |
| Ethnicity, n (%) Latino | 2 (11.1) | 2 (8.3) | 0.1 | 0.762 |
| Smoking Behaviors, n (%) | 0 (0.0) | 10 (41.7) | 9.8 | 0.002 |
| SMFQ Total Score | 3.8 (3.1) | 6.2 (5.4) | 1.6 | 0.110 |
| AUDIT-C | 0.0 (0.0) | 8.0 (1.0) | 8.0 | <.001 |

### Extraction of neuron-enriched exosomes

Quality and quantity of total exosomes extracted from plasma samples were analyzed by different methods. Ponceau S staining of the different fractions after size-exclusion chromatography showed a high purity of exosomal fractions with almost no contamination by plasma proteins (Figure 1A). Most plasma proteins were isolated with late protein fractions #7-10. Western blot analysis demonstrated that only exosomal fractions contain exosome-specific protein markers CD63 and TSG101 and, at the same time, showed no contamination with plasma albumin (Figure 1B). Interestingly, the neuronal marker L1CAM (CD171) was revealed in exosomal fractions but not in the fractions of the plasma proteins. TEM analysis demonstrated that the exosomal fractions contain vesicles with the shape and size corresponding to exosomes (Figure 1C). Nanoparticle tracking analysis showed average concentration of exosomes in plasma samples 3.1±0.8×10^11^ (particles/ml) (Figure 2A). After extraction from 0.5 ml of plasma sample, the exosomal fraction had a volume 1.5 ml with concentration 9.3±2.6×10^10^ (particles/ml). Hence, we were able to extract and purify ∼90% of plasma exosomes. After precipitation with anti-L1CAM (CD171) Abs, neuron-enriched exosomes were eluted in the 0.2 ml of elution buffer and demonstrated concentration 2.6±0.7×10^9^ (particles/ml). This result means that the neuron-enriched fraction of exosomes accounted for 0.04% of the total number of exosomes isolated from plasma. Exosome precipitation with the resin beads only (negative control) demonstrated that, in the absence of anti-L1CAM(CD171) Abs, concentration of precipitated exosomes was significantly lower (1.6±1.1×10^6^ particles/ml) (Figure 2A). Western blot analysis confirmed that while neuron-enriched and non-neuron exosomes expressed comparable levels of exosomal markers CD63 and TSG101, only neuron-enriched exosomes have a distinguishable level of L1CAM (CD171) protein expression (Figure 2B).

**Figure 1.**
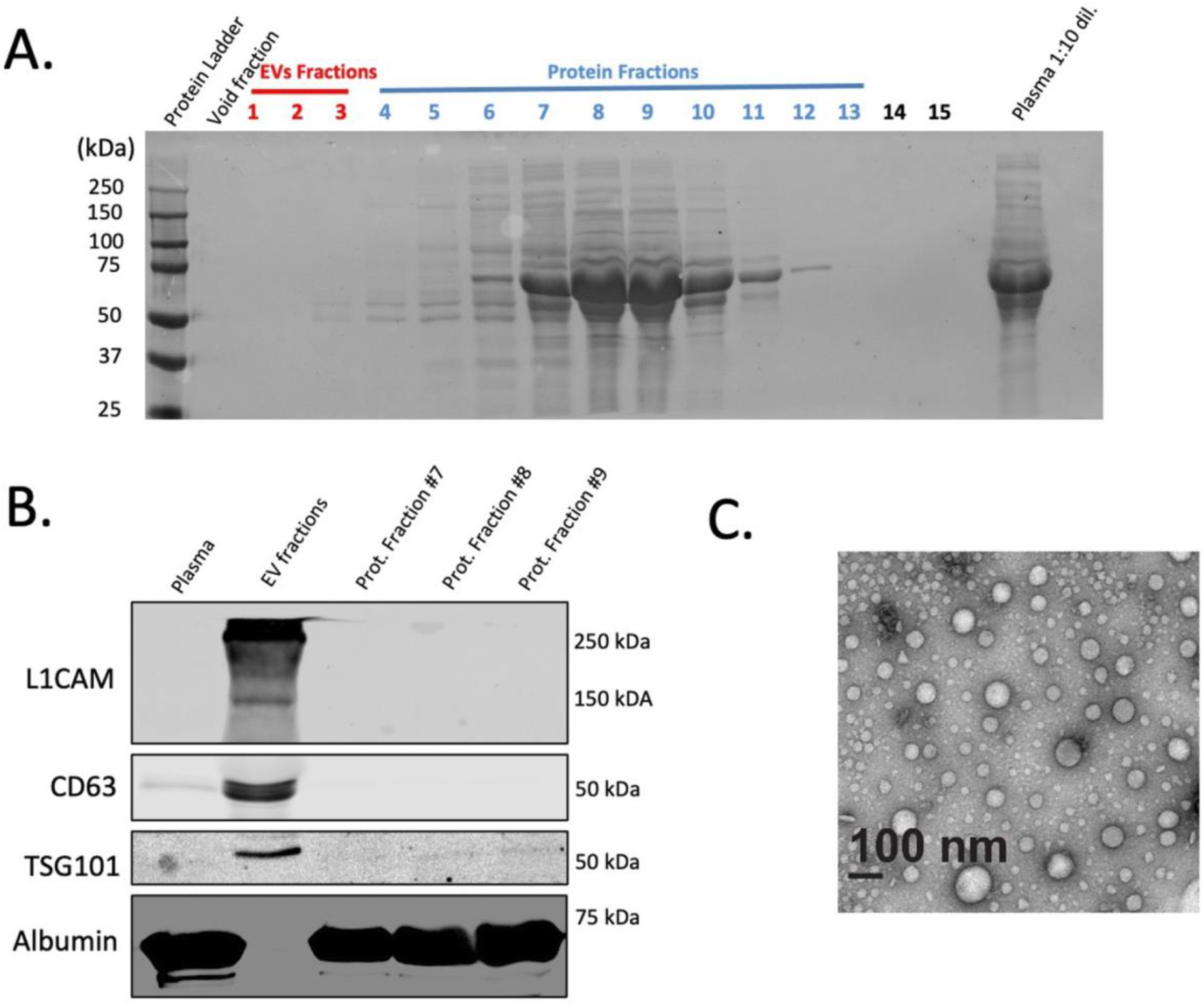
Extraction and characterization of exosomes from patient’s plasma samples. **(A)** Ponceau S staining of the different plasma fractions after extraction of exosomes by size-exclusion chromatography. **(B)** Western blot analysis of exosome-specific protein markers. Equal amount of total protein was loaded for native plasma, combined exosome fractions, and Protein fractions #7-9 (as shown in Figure 1A). **(C)** Electron microscopy analysis for the extracted exosomes.

**Figure 2.**
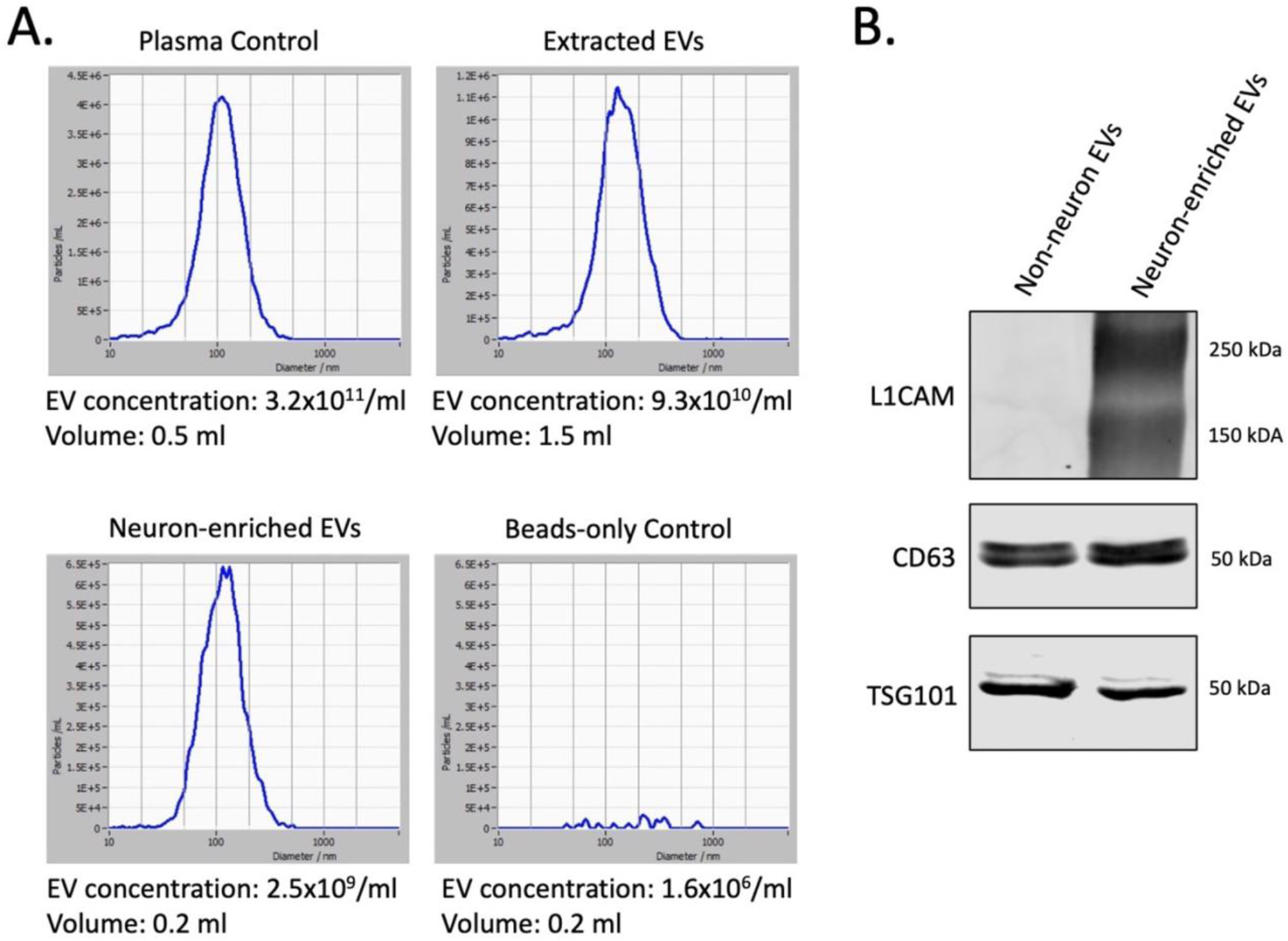
Precipitation and characterization of neuron-derived exosomes. **(A)** Nanoparticle tracking analysis of exosomes in the native plasma (Plasma Control), after extraction from plasma by the size-exclusion chromatography (Extracted EVs), after precipitation with Anti-L1CAM(CD171) Ab (Neuron-enriched EVs), and the beads-only control. **(B)** Analysis of exosome-specific markers and L1CAM(CD171) protein expression in neuron-derived exosomes and non-neuronal exosomes (supernatant). Equal amount of total protein from both samples were used for Western blot analysis.

### Analysis of miRNAs from neuron-enriched exosomes

The group-based microarray analysis identified 104 miRNAs, 6 of which showed the most stable expression and were identified as a potential normalization controls: miR-126-5p, miR-133a-3p, miR-139-5p, miR-197-5p, miR-328-3p, miR-199a-5p. Analysis of these miRNAs in all individual samples revealed that the most stable expression was demonstrated by miR-133a-3p and miR-197-5p (Figure S1). For further qPCR data analysis, miR-133a-3p was used as a normalization control. Out of 104 identified miRNAs, 22 showed at least a 50-fold difference in expression between heavy drinkers and controls. These 22 miRNAs were followed up with individual-level measurement via qPCR. One miRNA, mir-423-3p, produced unreliable qPCR measures and was removed from consideration. Analysis of individual samples showed no statistically significant difference between alcohol consumption and control groups for 18 out of the remaining 21 miRNAs (Figure S2). As presented in Table 2 and Figure 3, 3 of 21 miRNAs (miR-30a-5p, miR-194-5p, and miR-339-3p) were significantly differentially expressed by alcohol consumption group after controlling for age, sex, smoking behaviors, current depression symptoms, and accounting for twin pair relatedness; however, only miR-30a-5p and miR-194-5p survived Bonferroni correction. Both miR-30a-5p and miR-194-5p exhibited large effect sizes, with partial eta squared values of 0.26 and 0.24, respectively. Although miR-339-3p did not meet Bonferroni corrected significance, this miRNA was associated with a medium effect size. Supplemental Table 1 presents unadjusted and covariate adjusted means and standard errors for the 21 miRNAs while Supplemental Table 2 provides linear model parameter estimates for AU group and covariates on expression levels for non-significant miRNAs (*p* > .05).

**Figure 3.**
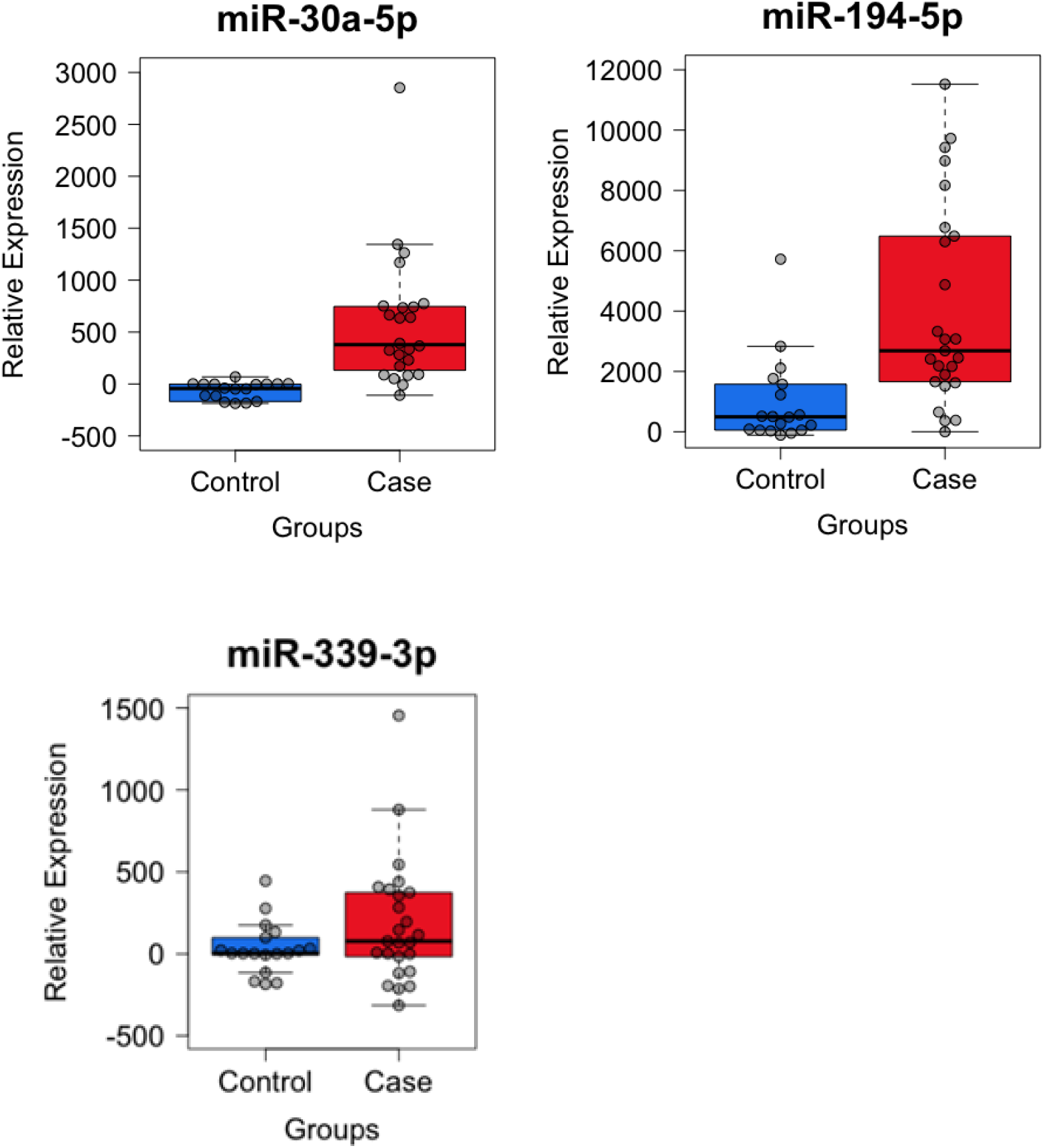
Comparison of Alcohol Use groups (controls versus cases) for neuron-enriched exosomal miRNA-30a-5p, miR-194-5p, and miR-339-3p expression.

**Table 2.**
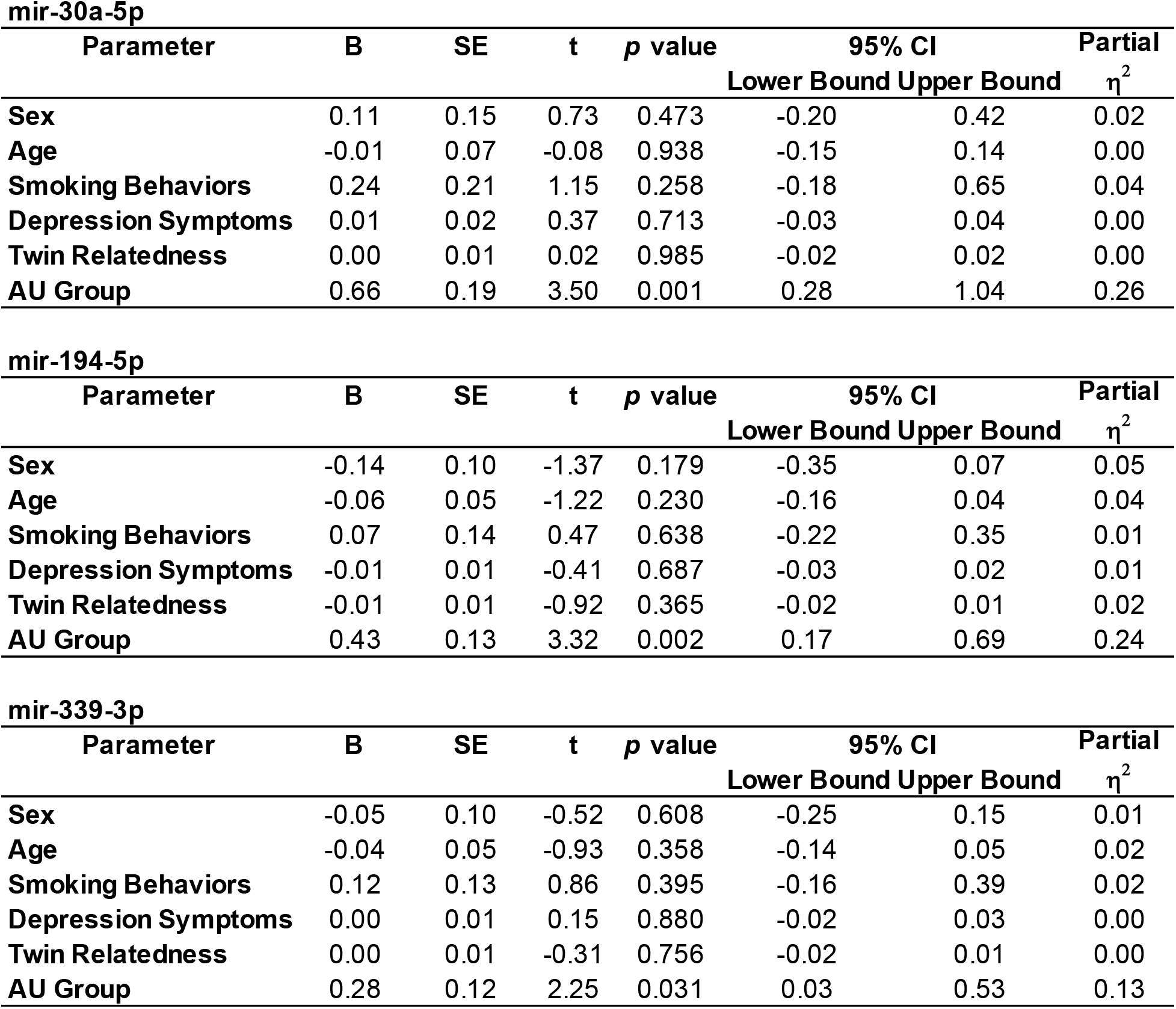
Linear model parameter estimates for Alcohol Use (AU) Group and covariates on expression levels of miRNA-30a-5p, miR-194-5p, and miR-339-3p.

| Parameter | B | SE | t | p value | 95% CI |  | Partial h<sup>2</sup> |
| --- | --- | --- | --- | --- | --- | --- | --- |
|  |  |  |  |  | Lower Bound | Upper Bound |  |
| **Sex** | 0.11 | 0.15 | 0.73 | 0.473 | -0.20 | 0.42 | 0.02 |
| **Age** | -0.01 | 0.07 | -0.08 | 0.938 | -0.15 | 0.14 | 0.00 |
| **Smoking Behaviors** | 0.24 | 0.21 | 1.15 | 0.258 | -0.18 | 0.65 | 0.04 |
| **Depression Symptoms** | 0.01 | 0.02 | 0.37 | 0.713 | -0.03 | 0.04 | 0.00 |
| **Twin Relatedness** | 0.00 | 0.01 | 0.02 | 0.985 | -0.02 | 0.02 | 0.00 |
| **AU Group** | 0.66 | 0.19 | 3.50 | 0.001 | 0.28 | 1.04 | 0.26 |

**mir-194-5p**
| Parameter | B | SE | t | p value | 95% CI |  | Partial h<sup>2</sup> |
| --- | --- | --- | --- | --- | --- | --- | --- |
|  |  |  |  |  | Lower Bound | Upper Bound |  |
| **Sex** | -0.14 | 0.10 | -1.37 | 0.179 | -0.35 | 0.07 | 0.05 |
| **Age** | -0.06 | 0.05 | -1.22 | 0.230 | -0.16 | 0.04 | 0.04 |
| **Smoking Behaviors** | 0.07 | 0.14 | 0.47 | 0.638 | -0.22 | 0.35 | 0.01 |
| **Depression Symptoms** | -0.01 | 0.01 | -0.41 | 0.687 | -0.03 | 0.02 | 0.01 |
| **Twin Relatedness** | -0.01 | 0.01 | -0.92 | 0.365 | -0.02 | 0.01 | 0.02 |
| **AU Group** | 0.43 | 0.13 | 3.32 | 0.002 | 0.17 | 0.69 | 0.24 |

**mir-339-3p**
| Parameter | B | SE | t | p value | 95% CI |  | Partial h<sup>2</sup> |
| --- | --- | --- | --- | --- | --- | --- | --- |
|  |  |  |  |  | Lower Bound | Upper Bound |  |
| **Sex** | -0.05 | 0.10 | -0.52 | 0.608 | -0.25 | 0.15 | 0.01 |
| **Age** | -0.04 | 0.05 | -0.93 | 0.358 | -0.14 | 0.05 | 0.02 |
| **Smoking Behaviors** | 0.12 | 0.13 | 0.86 | 0.395 | -0.16 | 0.39 | 0.02 |
| **Depression Symptoms** | 0.00 | 0.01 | 0.15 | 0.880 | -0.02 | 0.03 | 0.00 |
| **Twin Relatedness** | 0.00 | 0.01 | -0.31 | 0.756 | -0.02 | 0.01 | 0.00 |
| **AU Group** | 0.28 | 0.12 | 2.25 | 0.031 | 0.03 | 0.53 | 0.13 |

### MiRNA-miRNA Interaction Network Inference

The miRNA-miRNA interaction network inferred by the miRsig tool for the alcohol consumption network is shown in Figure 4A. This network consists of 16 miRNAs and 77 edges arranged in two connected clusters. None of the miRNAs in this predicted network were differentially expressed (DE) by case status. Let-7i-5p miRNA was found to be the highest-degree node in this network. For a sensitivity analysis, we recreated the miRNA-miRNA interaction network with only unrelated individuals by dropping one twin from each of the six twin pairs. No significant difference in the general network structure was observed after dropping these twins (Figure 4B). Let-7i-5p was still ranked as the highest-degree node in the network, and the two networks shared 88% of their edges (61/69).

**Figure 4.**
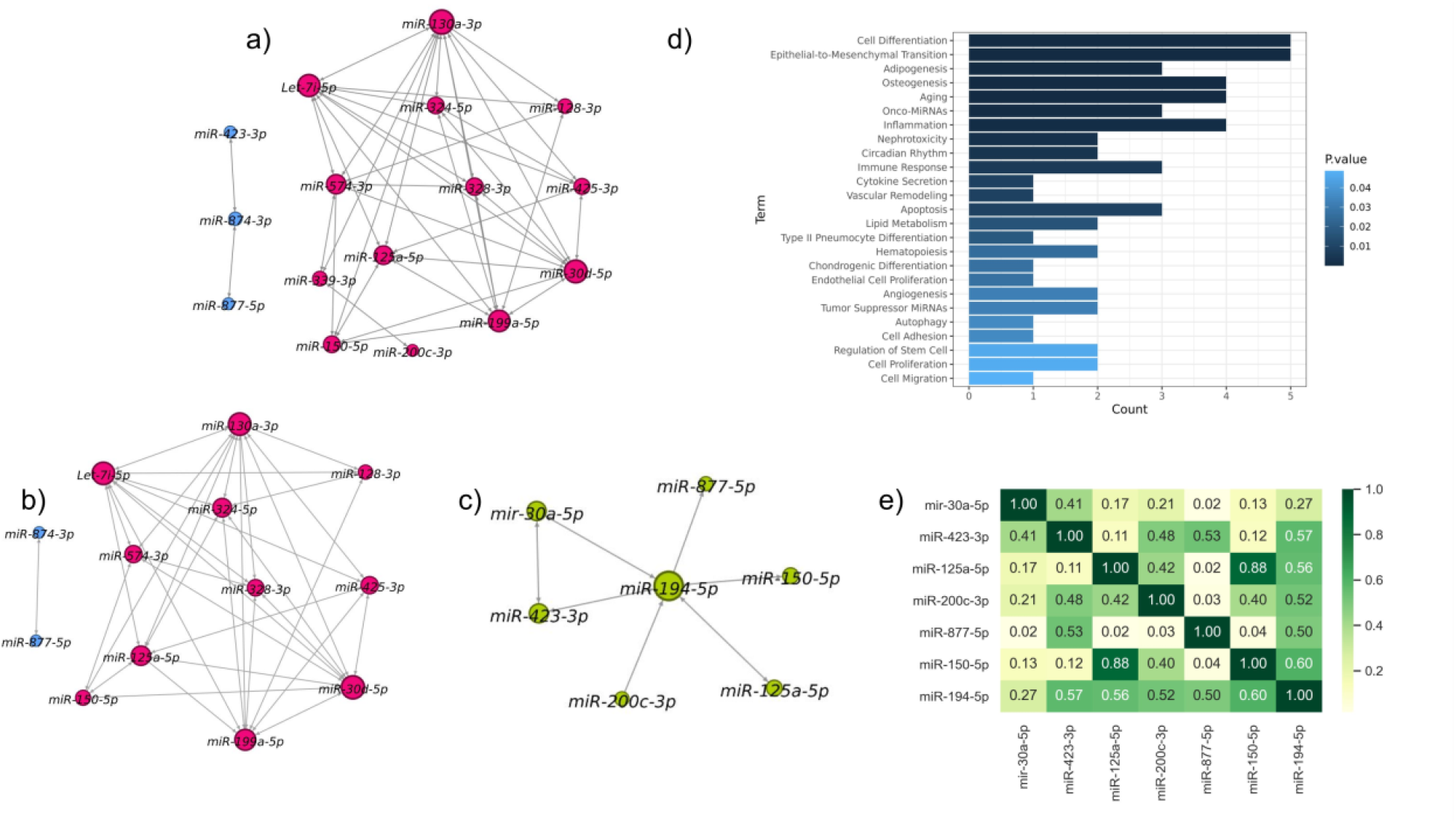
MiRNA-miRNA interaction network inference. **(A)** Inferred miRNA-miRNA interaction network. This network consists of 16 miRNAs and 77 edges and two clusters. **(B)** Inferred miRNA-miRNA interaction network after dropping the twin participants. This network consists of 13 miRNAs and 69 edges and two clusters. **(C)** Differentially expressed miRNAs and their direct regulators consist of seven miRNAs. **(D)** Functional analysis of the seven miRNAs from (C) comprising two differentially expressed miRNAs and their direct regulators. 25 biological functions are statistically significant. **(E)** Correlation between differentially expressed miRNAs and their direct regulators. miR-194-5p is highly correlated with the other miRNAs in this cluster.

### miRNAs interacting with DE miRNAs

Because none of the differentially expressed miRNAs from the regression models appeared in our initial miRNA-miRNA interaction network, we built another network with differentially expressed miRNAs and their direct regulators (neighbors in the network) by rerunning the miRsig pipeline with a reduced consensus cutoff (original cutoff 0.8, new cutoff 0.7) so that all the differentially expressed miRNAs would be included. This second network consisted of seven miRNAs thought to work together (Figure 4C). A functional analysis identified 25 biological functions significantly associated with these miRNAs (Figure 4D). miR-194-5p was ranked the highest-degree node in this network and its expression was highly correlated with the other miRNAs in this cluster (Figure 4E).

## Discussion

We used a commercially available microarray platform to identify neuron-enriched exosomal miRNAs differentially expressed by self-reported alcohol consumption rate in a healthy sample of young people. Follow up molecular studies validated the microarray results, and multivariate regression models controlling for demographics and clinical features identified mi R-30a-5p and miR-194-5p as significantly differentially expressed between cases and controls. MiR-30a-5p is one of several microRNAs that can regulate the expression of the brain-derived neurotrophic factor (*BDNF*) gene. Indeed, in a mouse model used to approximate binge drinking, overexpression of miR-30a-5p in the prefrontal cortex was associated with downregulation of *BDNF* expression and increased alcohol consumption^41^. The replication and observed effect size suggest that miR-30a-5p expression differences associated with behaviors like alcohol use may follow conserved biological networks, be replicable across multiple species, and provide insight into interindividual differences in health outcomes. MiR-194-5p also was significantly differentially overexpressed in young people reporting high alcohol consumption rates, and this miRNA was the highest-degree node in the miRNA network, correlating highly with the other miRNAs in this cluster. Moreover, the effect size for these two miRNA was substantial, suggesting a robust association. A functional analysis of the miRNAs belonging to this cluster implicated 25 biological functions, with the most significant terms related to cell differentiation, inflammation, aging, epithelial-mesenshymal transition, osteogenesis, apoptosis, immune response, and onco-miRNAs. Overall, our research suggests that alcohol consumption, even in young people, may affect the expression of circulating exosomal microRNAs and cellular regulatory mechanisms. Indeed, it is possible that dysregulated alcohol consumption may affect the production of critical factors and proteins like BDNF throughout the body, leading to a neurobiological landscape with elevated risks for psychiatric and substance use disorders.

By examining the microRNA profiles of circulating exosomes in an adolescent and young adult cohort, this study addresses several important gaps in the literature. First, it provides evidence that ample exosomes circulate in the peripheral blood of healthy young people to justify pursuing additional work to characterize not only the roles of exosomes during adolescence but also in-depth fundamental analyses to evaluate the variability of exosome number, size, type, and cargo in healthy, typically developing individuals. More research is needed to understand how biological factors (e.g., sex), genetic variation, behavior (e.g., substance use), health status (e.g., HIV status), environmental factors (e.g., exposure to pollution) and demographic factors (e.g., race, age) impact exosome cargo, quantity, and destinations. Such foundational knowledge would bolster future translational work to assess the utility of exosomes in clinical applications like screening tests, prognostic indicators, and precision medicine initiatives.

Second, this study indicates that high throughput omics technology like next-generation sequencing platforms to identify and quantify both known and novel RNA cargo may be feasible approaches for future exosomal miRNA research in adolescents and young people based on the total amount of exosomal RNA available in a modest plasma sample. In our cohort, we were able to extract 181.2 ± 42.1 ng of total RNA from the neuron-enriched exosomes of each blood sample. Given that this small amount of RNA cannot be applied for the next-generation sequencing analysis, we opted to use a commercially available qPCR serum/plasma-focused panel to measure the microRNAs expression. This platform is affordable and generates modest amounts of count data that do not require specialized data wrangling skills or computer clusters to manage. However, a qPCR method is not an ideal approach for quantifying novel or unexpected RNAs because qPCR and microarrays only measure RNAs represented by in the probe set. Thus, any microRNA present in a sample but unaccounted for in the microarray probe set will go unmeasured. Other work in adult populations suggest that additional noncoding RNA species may be present and in need of characterization.

Third, this study provides evidence that precipitation with Anti-L1CAM Abs is an effective and reliable method of enrichment of L1CAM-associated exosomes from the human plasma samples. Previously, Norman et al. (2021) expressed concern that most of the L1CAM observed in exosome studies was soluble protein not associated with exosomes in CSF or plasma^69^. While we cannot speak to CSF, we demonstrated that L1CAM protein is located mainly in the exosome fraction and not in the fraction of free plasma proteins (Figure 1). Our work cannot give a clear explanation of this discrepancy. The probable reason is different approaches to exosome isolation from plasma and/or blood sample handling. Possibly, damage to exosomes during certain extraction methods or improper storage of plasma samples could lead to the release of L1CAM protein from exosomes into the fraction of free plasma proteins. Additionally, although we do not have exosome tracking data to confirm our L1CAM-enriched sample was primarily of neuronal origin, we are encouraged by both our own results and other findings in the literature that have identified miRNAs and proteins associated with brain disorders and brain cell activity. Collectively, these reports support the hypothesis that L1CAM is indicative of neuronal origin and that selecting exosomes based on L1CAM association enriches the analytic sample for exosomes of brain cell origin. Regardless of the accuracy of the L1CAM hypothesis, we strongly encourage future work characterizing L1CAM-associated exosomes and suggest that even if L1CAM is not a definitive marker of neuronal origin, it has been an informative marker with relation to several mood and neuro-related disorders (e.g., Alzheimer’s, Parkinsons, amyotrophic lateral sclerosis [ALS], brain injury, bipolar depression) and that this utility remains regardless of L1CAM’s specificity to neurons/neural tissue^38, 70–73^.

This study is not without limitations. First, despite being one of the largest studies of healthy young people, the sample size is modest and relatively homogeneous in genetic background and socio-demographic variables like age and race. A larger, more diverse cohort is needed to explore possible age, sex, and race/ethnic related effects, and twin studies may be needed to achieve accurate estimates of the magnitude of genetic and environmental influences on exosome quantity and cargo composition. Second, while the two pronged measurement strategy of group-based microarray followed by quantitative PCR minimized financial risk, it limited data resolution; individual-level microRNA measures were collected only for the microRNAs that exhibited strong group differences, meaning that microRNAs of modest effect size or that only showed differential expression in a subset of cases were likely overlooked. Third, our analyses included almost no individuals who engaged in low-to-moderate alcohol consumption, which limited our ability to perform quantitative analyses (as opposed to categorical group analyses). A larger cohort representing a continuous distribution of drinking rates (none, low, moderate, and heavy) would increase statistical power and enable more sophisticated statistical modeling techniques. Collectively, the associations observed between neuron-enriched exosomal miRNAs and alcohol use suggests that high rates of alcohol consumption during the adolescent/young adult years may impact brain maturation and functioning by modulating miRNA expression.

## Acknowledgements and Disclosures

The AYATS parent study was supported by NIH/NIMH R01MH101518 to the last author (RRN). AYATS received IRB approval from the Virginia Commonwealth University IRB (protocol #HM20022942). This study and VY were supported by funds from the VCU Alcohol Research Center (P50AA022537) and from the VCU Department of Psychiatry to the last author (RRN). A National Center for Advancing Translational Sciences grant (UL1TR000058) supported resources used to conduct AYATS. PR and PG were supported by the Commonwealth Health Research Board (CHRB #236-06-23) and NSF (CBET-1802588) awarded to PG. VY also was supported by the internal fund of the VCU Massey Cancer Center. Services and products in support of the research project were generated by the VCU Massey Cancer Center Microscopy Core Shared Resource, supported, in part, with funding from NIH-NCI Cancer Center Support Grant P30 CA016059. None of the authors report potential conflicts of interest. The data and code related to this publication have been deposited to the Open Science Framework (https://osf.io/gaw79).

## Supplement

**Supplemental Table 1.**
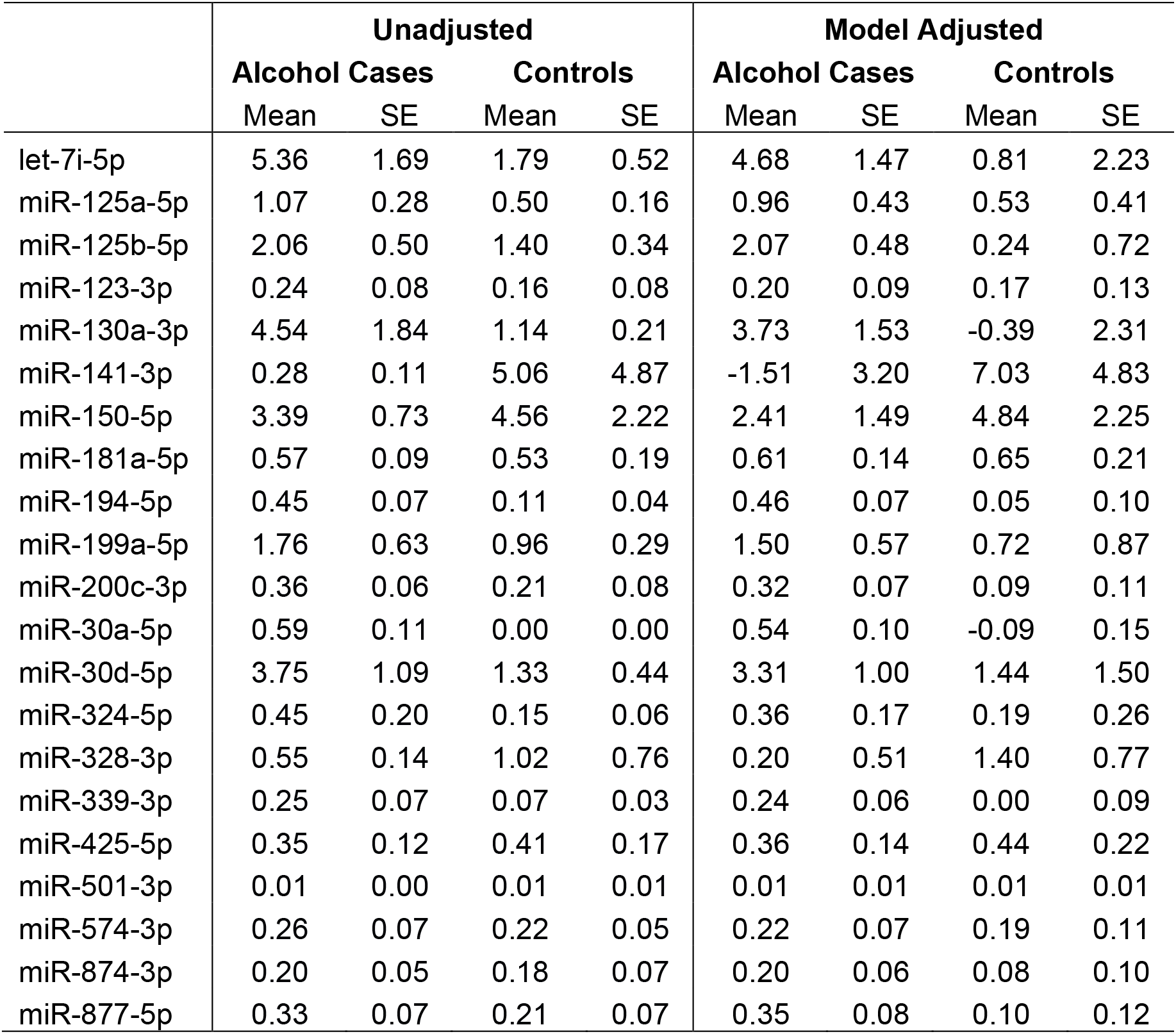
Unadjusted and covariate adjusted means and standard errors.

**Supplemental Table 2.**
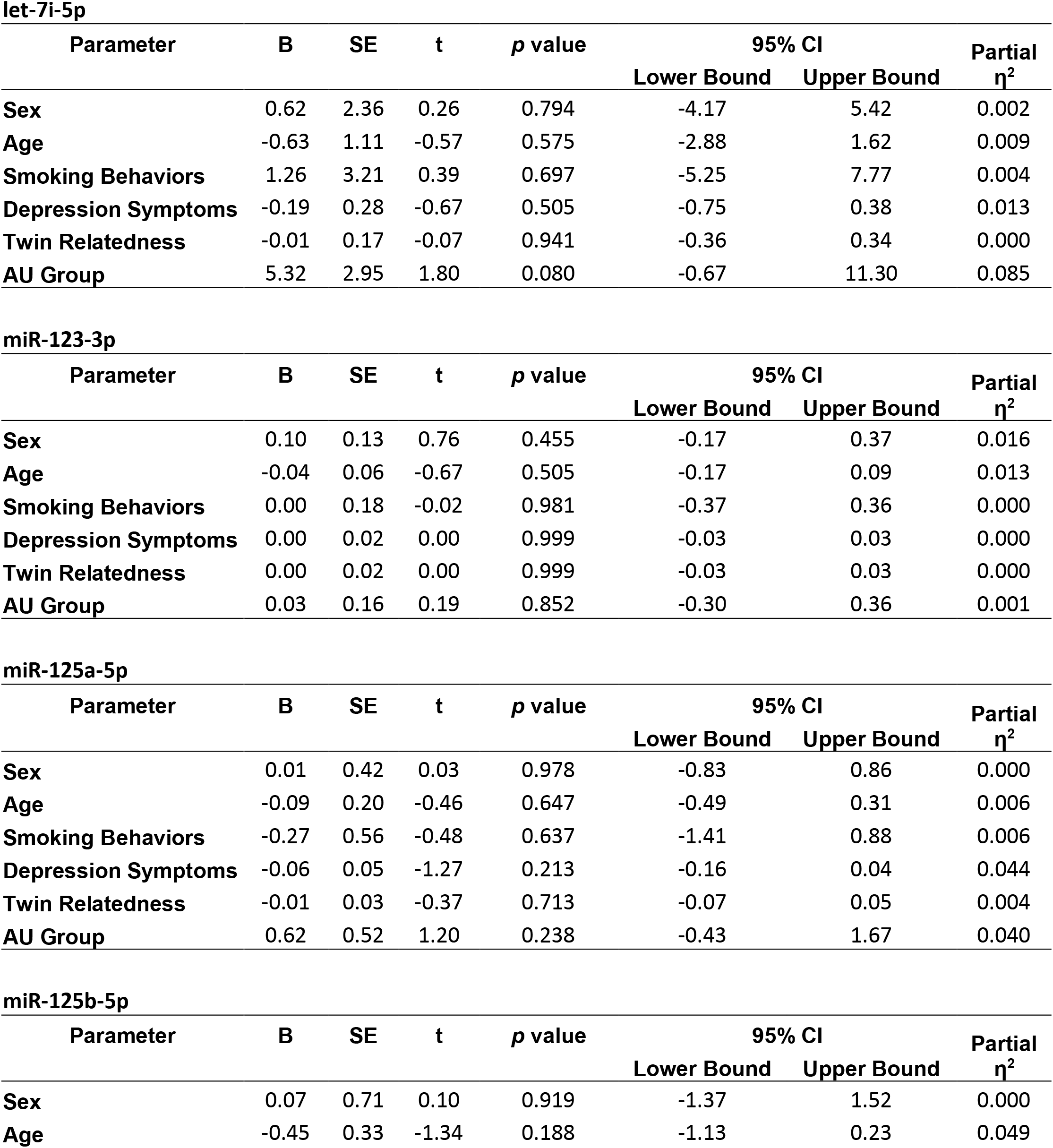
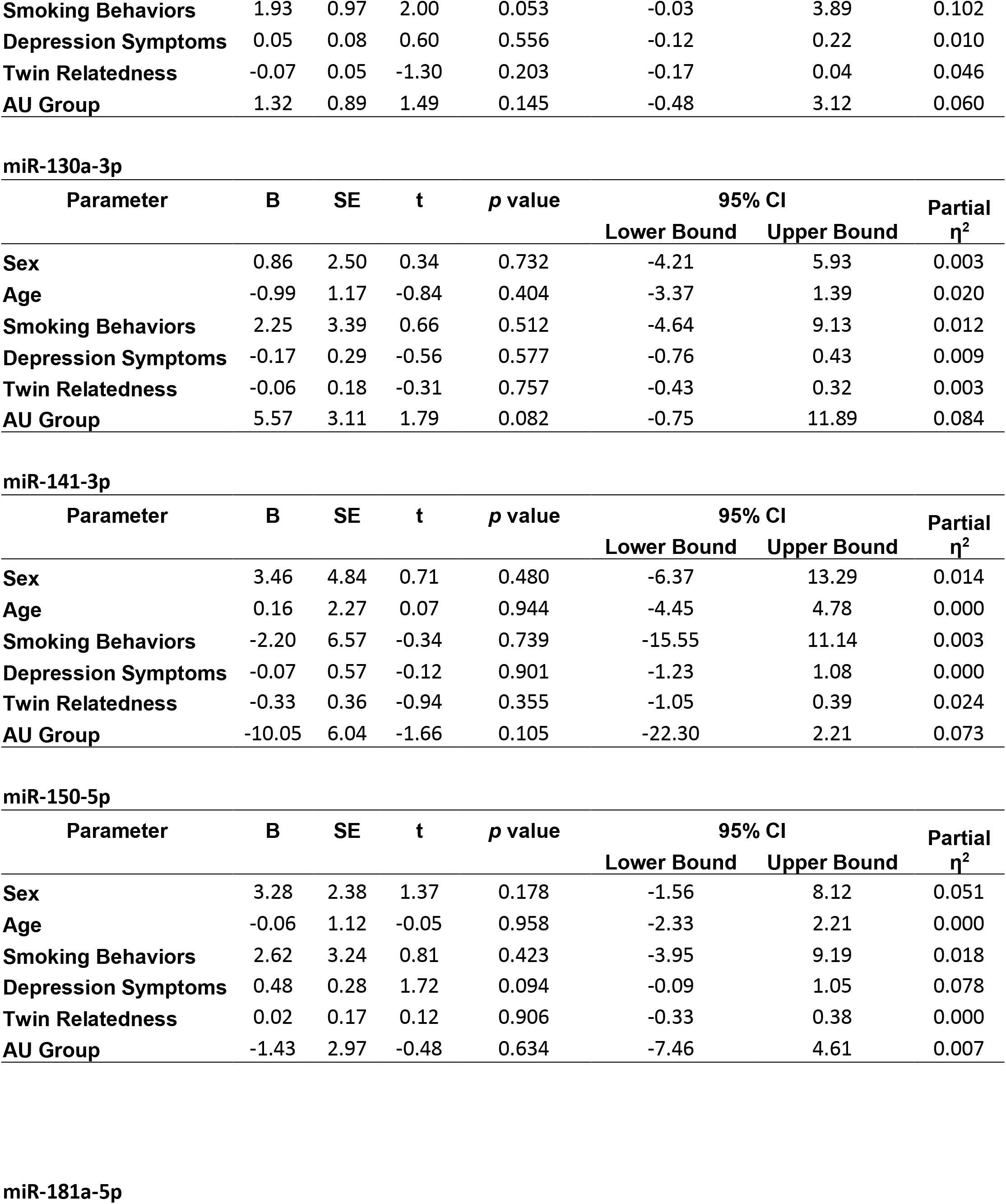
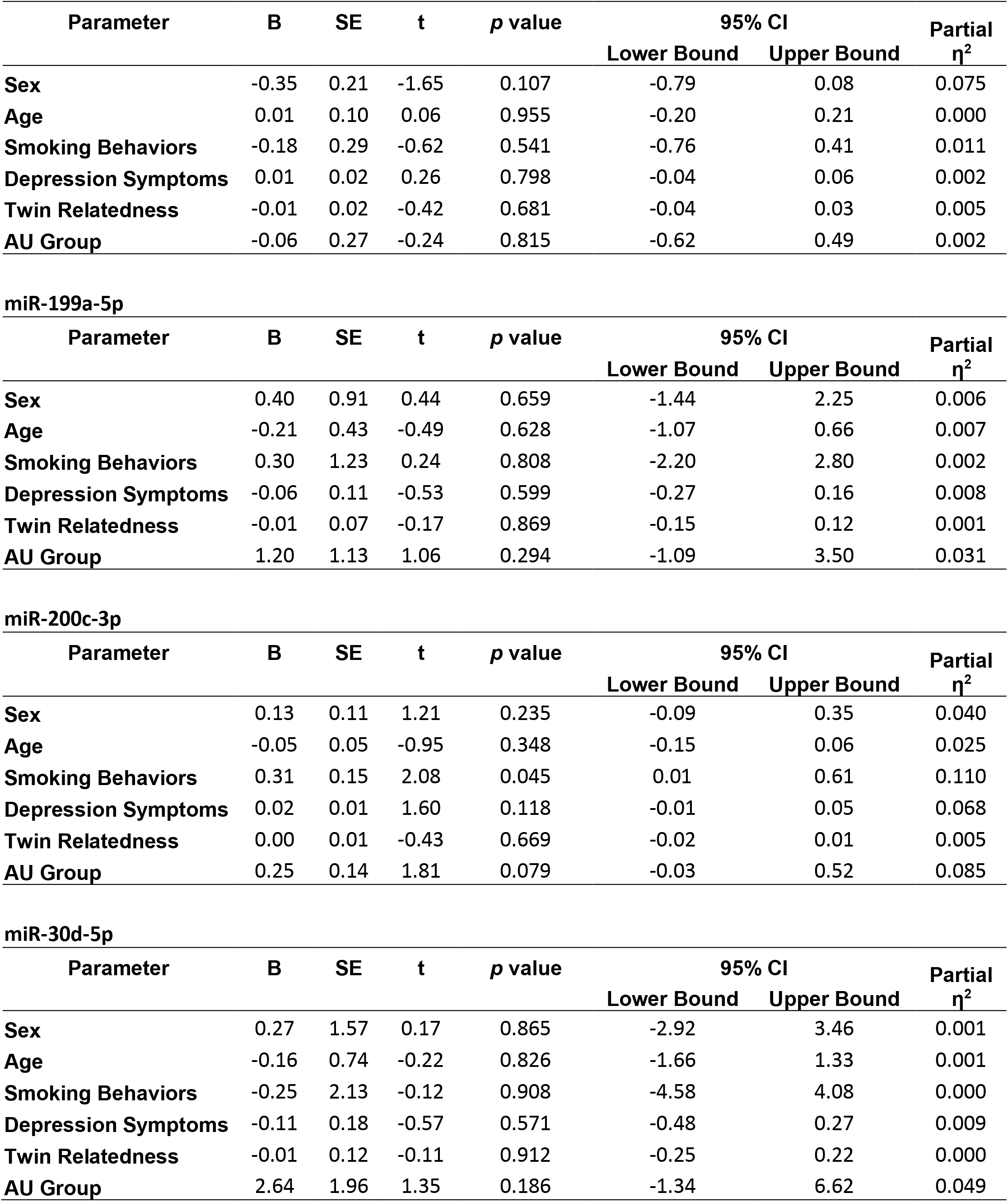
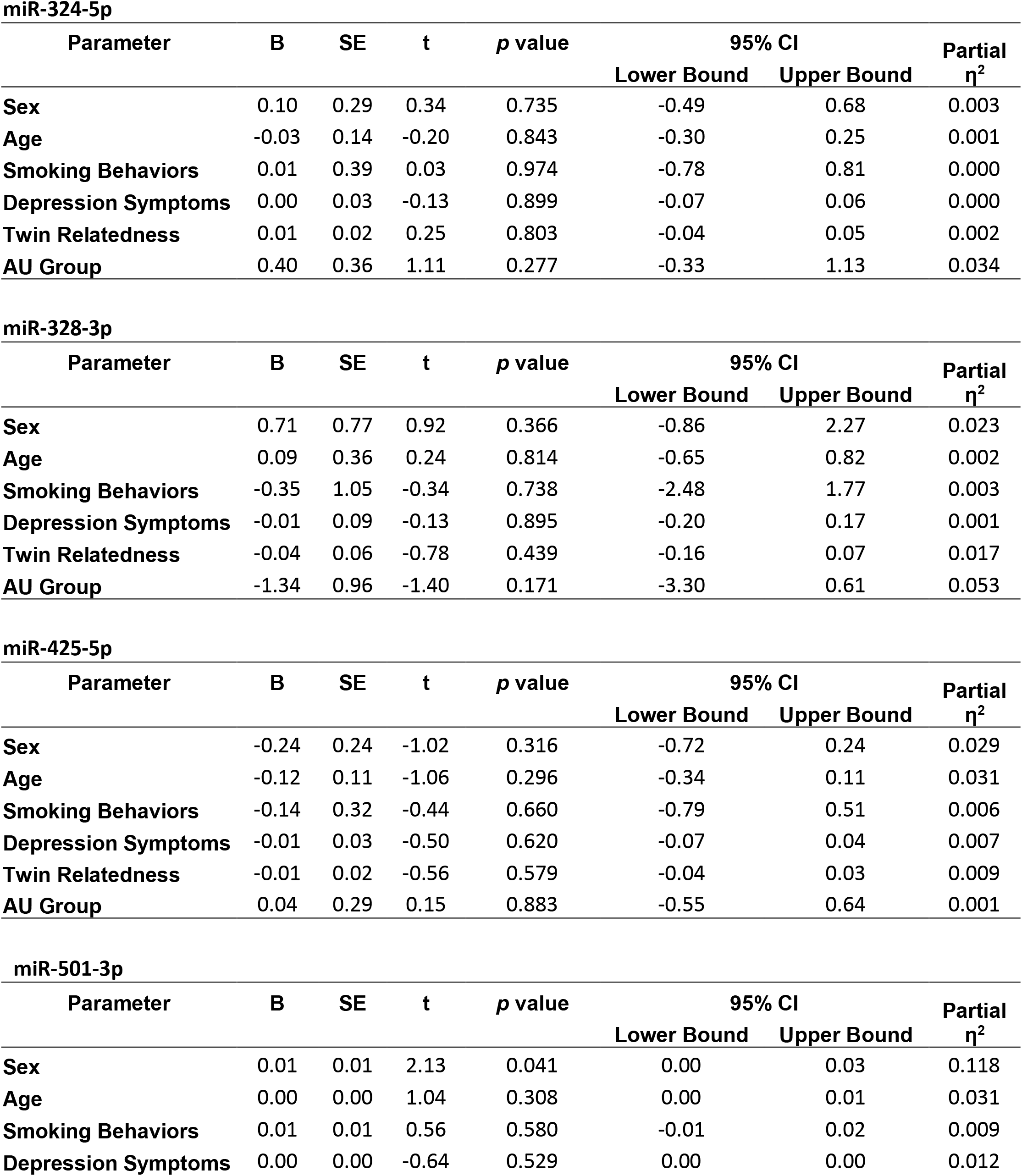
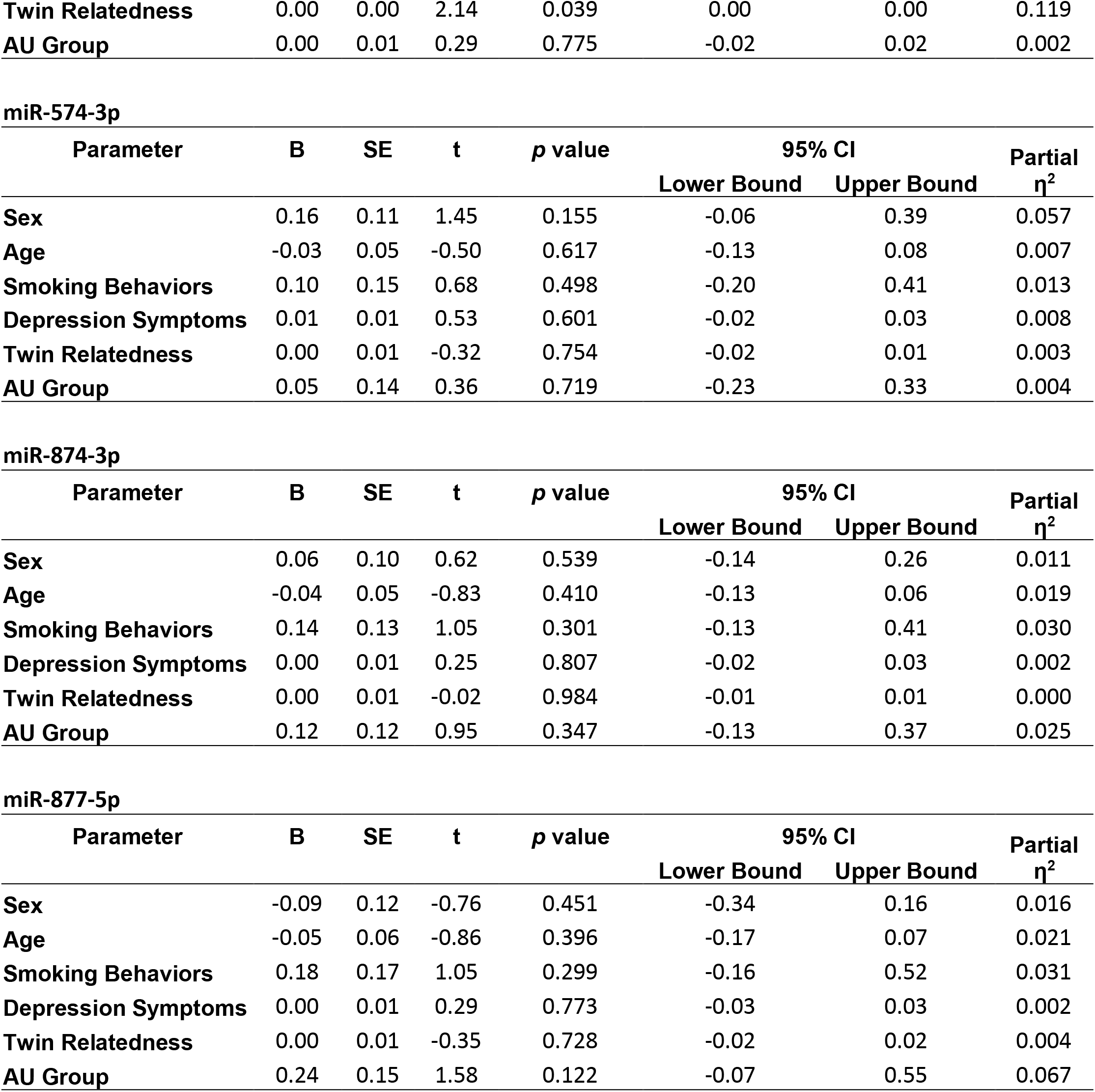
Linear model parameter estimates for Alcohol Use (AU) Group and covariates on expression levels for non-significant miRNAs.

**Figure S1.**
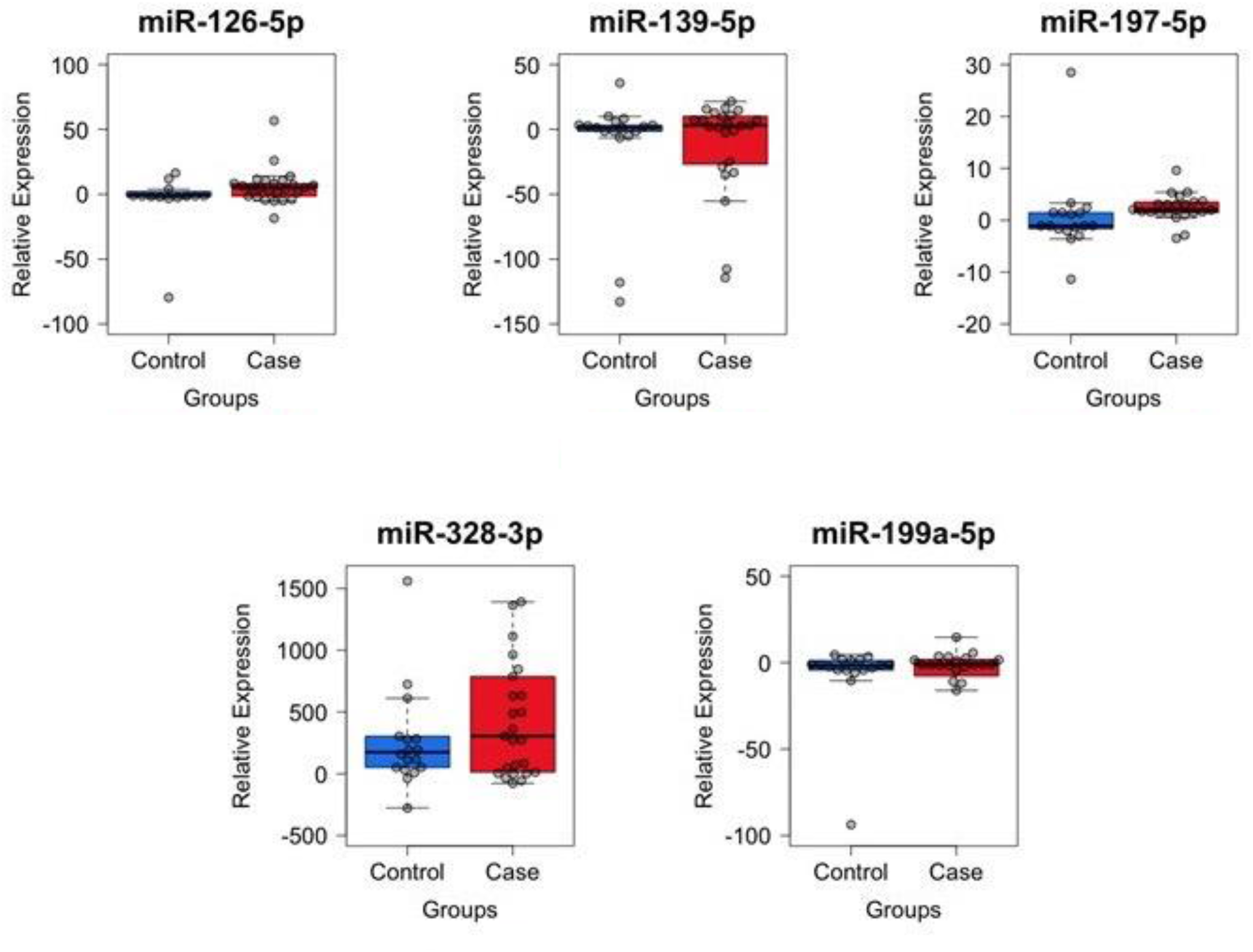
Potential internal controls normalized by miR-133a-3p

**Figure S2.**
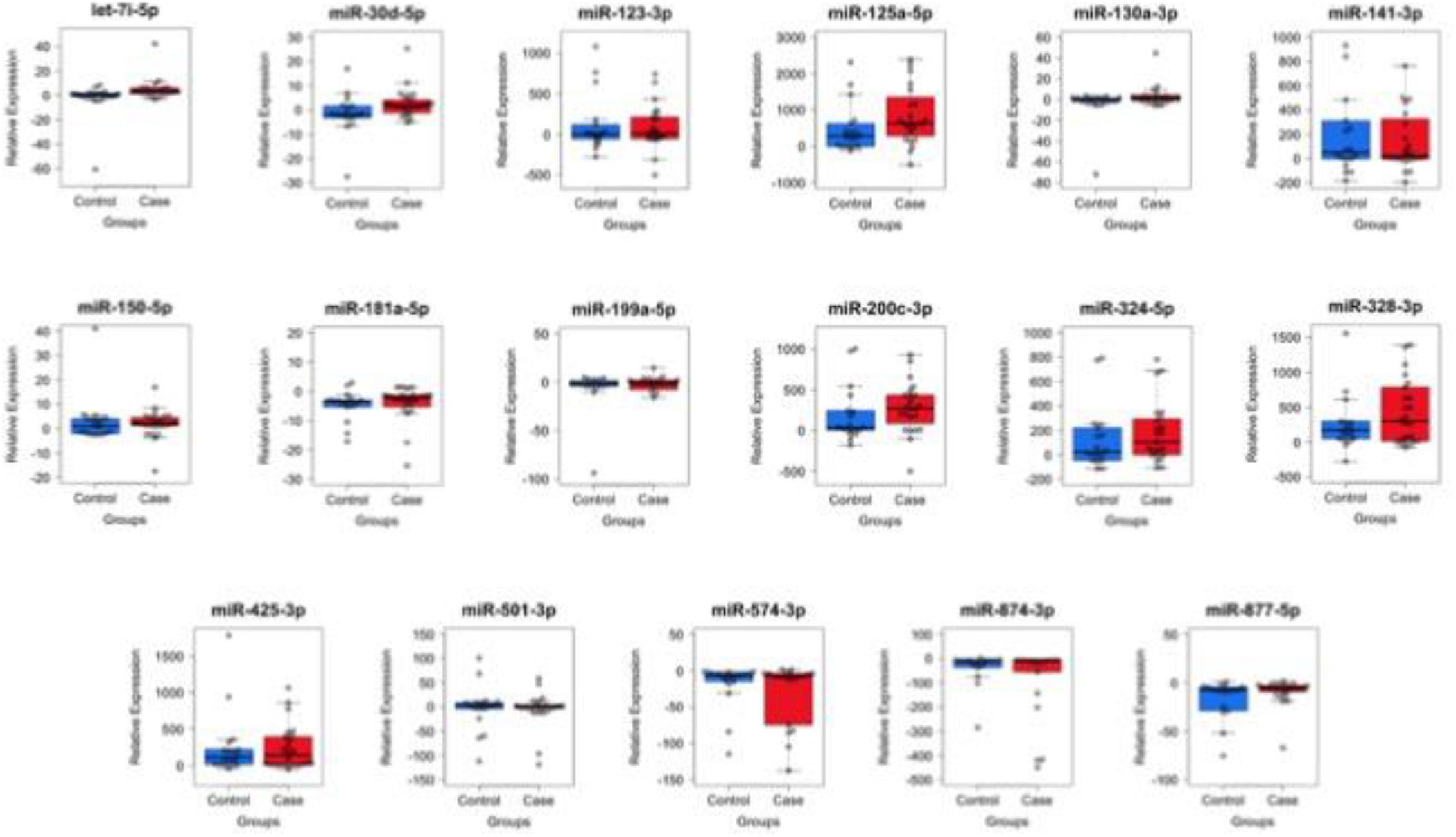
qPCR analysis of individual samples from alcohol consumption (Case) and control groups.

## Notes

https://osf.io/gaw79

### Competing Interest Statement

The authors have declared no competing interest.

