## Supplemental_Tables_and_Figures for "Neuron Enriched Exosomal MicroRNA Expression Profiles as a Marker of Early Life Alcohol Consumption"

**Supplement**

**Supplemental Table 1. Unadjusted and covariate adjusted means and standard errors.**

| | **Unadjusted** | | | | **Model Adjusted** | | | |
|---|---|---|---|---|---|---|---|---|
| | **Alcohol Cases** | | **Controls** | | **Alcohol Cases** | | **Controls** | |
| | Mean | SE | Mean | SE | Mean | SE | Mean | SE |
| let-7i-5p | 5.36 | 1.69 | 1.79 | 0.52 | 4.68 | 1.47 | 0.81 | 2.23 |
| miR-125a-5p | 1.07 | 0.28 | 0.50 | 0.16 | 0.96 | 0.43 | 0.53 | 0.41 |
| miR-125b-5p | 2.06 | 0.50 | 1.40 | 0.34 | 2.07 | 0.48 | 0.24 | 0.72 |
| miR-123-3p | 0.24 | 0.08 | 0.16 | 0.08 | 0.20 | 0.09 | 0.17 | 0.13 |
| miR-130a-3p | 4.54 | 1.84 | 1.14 | 0.21 | 3.73 | 1.53 | -0.39 | 2.31 |
| miR-141-3p | 0.28 | 0.11 | 5.06 | 4.87 | -1.51 | 3.20 | 7.03 | 4.83 |
| miR-150-5p | 3.39 | 0.73 | 4.56 | 2.22 | 2.41 | 1.49 | 4.84 | 2.25 |
| miR-181a-5p | 0.57 | 0.09 | 0.53 | 0.19 | 0.61 | 0.14 | 0.65 | 0.21 |
| miR-194-5p | 0.45 | 0.07 | 0.11 | 0.04 | 0.46 | 0.07 | 0.05 | 0.10 |
| miR-199a-5p | 1.76 | 0.63 | 0.96 | 0.29 | 1.50 | 0.57 | 0.72 | 0.87 |
| miR-200c-3p | 0.36 | 0.06 | 0.21 | 0.08 | 0.32 | 0.07 | 0.09 | 0.11 |
| miR-30a-5p | 0.59 | 0.11 | 0.00 | 0.00 | 0.54 | 0.10 | -0.09 | 0.15 |
| miR-30d-5p | 3.75 | 1.09 | 1.33 | 0.44 | 3.31 | 1.00 | 1.44 | 1.50 |
| miR-324-5p | 0.45 | 0.20 | 0.15 | 0.06 | 0.36 | 0.17 | 0.19 | 0.26 |
| miR-328-3p | 0.55 | 0.14 | 1.02 | 0.76 | 0.20 | 0.51 | 1.40 | 0.77 |
| miR-339-3p | 0.25 | 0.07 | 0.07 | 0.03 | 0.24 | 0.06 | 0.00 | 0.09 |
| miR-425-5p | 0.35 | 0.12 | 0.41 | 0.17 | 0.36 | 0.14 | 0.44 | 0.22 |
| miR-501-3p | 0.01 | 0.00 | 0.01 | 0.01 | 0.01 | 0.01 | 0.01 | 0.01 |
| miR-574-3p | 0.26 | 0.07 | 0.22 | 0.05 | 0.22 | 0.07 | 0.19 | 0.11 |
| miR-874-3p | 0.20 | 0.05 | 0.18 | 0.07 | 0.20 | 0.06 | 0.08 | 0.10 |
| miR-877-5p | 0.33 | 0.07 | 0.21 | 0.07 | 0.35 | 0.08 | 0.10 | 0.12 |

**Supplemental Table 2. Linear model parameter estimates for Alcohol Use (AU) Group and covariates on expression levels for non-significant miRNAs.**

**let-7i-5p**

| Parameter | B | SE | t | *p* value | 95% CI | | Partial $\eta^2$ |
|---|---|---|---|---|---|---|---|
| | | | | | Lower Bound | Upper Bound | |
| **Sex** | 0.62 | 2.36 | 0.26 | 0.794 | -4.17 | 5.42 | 0.002 |
| **Age** | -0.63 | 1.11 | -0.57 | 0.575 | -2.88 | 1.62 | 0.009 |
| **Smoking Behaviors** | 1.26 | 3.21 | 0.39 | 0.697 | -5.25 | 7.77 | 0.004 |
| **Depression Symptoms** | -0.19 | 0.28 | -0.67 | 0.505 | -0.75 | 0.38 | 0.013 |
| **Twin Relatedness** | -0.01 | 0.17 | -0.07 | 0.941 | -0.36 | 0.34 | 0.000 |
| **AU Group** | 5.32 | 2.95 | 1.80 | 0.080 | -0.67 | 11.30 | 0.085 |

**miR-123-3p**

| Parameter | B | SE | t | *p* value | 95% CI | | Partial $\eta^2$ |
|---|---|---|---|---|---|---|---|
| | | | | | Lower Bound | Upper Bound | |
| **Sex** | 0.10 | 0.13 | 0.76 | 0.455 | -0.17 | 0.37 | 0.016 |
| **Age** | -0.04 | 0.06 | -0.67 | 0.505 | -0.17 | 0.09 | 0.013 |
| **Smoking Behaviors** | 0.00 | 0.18 | -0.02 | 0.981 | -0.37 | 0.36 | 0.000 |
| **Depression Symptoms** | 0.00 | 0.02 | 0.00 | 0.999 | -0.03 | 0.03 | 0.000 |
| **Twin Relatedness** | 0.00 | 0.02 | 0.00 | 0.999 | -0.03 | 0.03 | 0.000 |
| **AU Group** | 0.03 | 0.16 | 0.19 | 0.852 | -0.30 | 0.36 | 0.001 |

**miR-125a-5p**

| Parameter | B | SE | t | *p* value | 95% CI | | Partial $\eta^2$ |
|---|---|---|---|---|---|---|---|
| | | | | | Lower Bound | Upper Bound | |
| **Sex** | 0.01 | 0.42 | 0.03 | 0.978 | -0.83 | 0.86 | 0.000 |
| **Age** | -0.09 | 0.20 | -0.46 | 0.647 | -0.49 | 0.31 | 0.006 |
| **Smoking Behaviors** | -0.27 | 0.56 | -0.48 | 0.637 | -1.41 | 0.88 | 0.006 |
| **Depression Symptoms** | -0.06 | 0.05 | -1.27 | 0.213 | -0.16 | 0.04 | 0.044 |
| **Twin Relatedness** | -0.01 | 0.03 | -0.37 | 0.713 | -0.07 | 0.05 | 0.004 |
| **AU Group** | 0.62 | 0.52 | 1.20 | 0.238 | -0.43 | 1.67 | 0.040 |

**miR-125b-5p**

| Parameter | B | SE | t | *p* value | 95% CI | | Partial $\eta^2$ |
|---|---|---|---|---|---|---|---|
| | | | | | Lower Bound | Upper Bound | |
| **Sex** | 0.07 | 0.71 | 0.10 | 0.919 | -1.37 | 1.52 | 0.000 |
| **Age** | -0.45 | 0.33 | -1.34 | 0.188 | -1.13 | 0.23 | 0.049 |

| | | | | | | | |
|---|---|---|---|---|---|---|---|
| **Smoking Behaviors** | 1.93 | 0.97 | 2.00 | 0.053 | -0.03 | 3.89 | 0.102 |
| **Depression Symptoms** | 0.05 | 0.08 | 0.60 | 0.556 | -0.12 | 0.22 | 0.010 |
| **Twin Relatedness** | -0.07 | 0.05 | -1.30 | 0.203 | -0.17 | 0.04 | 0.046 |
| **AU Group** | 1.32 | 0.89 | 1.49 | 0.145 | -0.48 | 3.12 | 0.060 |

**miR-130a-3p**

| Parameter | B | SE | t | *p* value | 95% CI | | Partial $\eta^2$ |
|---|---|---|---|---|---|---|---|
| | | | | | Lower Bound | Upper Bound | |
| **Sex** | 0.86 | 2.50 | 0.34 | 0.732 | -4.21 | 5.93 | 0.003 |
| **Age** | -0.99 | 1.17 | -0.84 | 0.404 | -3.37 | 1.39 | 0.020 |
| **Smoking Behaviors** | 2.25 | 3.39 | 0.66 | 0.512 | -4.64 | 9.13 | 0.012 |
| **Depression Symptoms** | -0.17 | 0.29 | -0.56 | 0.577 | -0.76 | 0.43 | 0.009 |
| **Twin Relatedness** | -0.06 | 0.18 | -0.31 | 0.757 | -0.43 | 0.32 | 0.003 |
| **AU Group** | 5.57 | 3.11 | 1.79 | 0.082 | -0.75 | 11.89 | 0.084 |

**miR-141-3p**

| Parameter | B | SE | t | *p* value | 95% CI | | Partial $\eta^2$ |
|---|---|---|---|---|---|---|---|
| | | | | | Lower Bound | Upper Bound | |
| **Sex** | 3.46 | 4.84 | 0.71 | 0.480 | -6.37 | 13.29 | 0.014 |
| **Age** | 0.16 | 2.27 | 0.07 | 0.944 | -4.45 | 4.78 | 0.000 |
| **Smoking Behaviors** | -2.20 | 6.57 | -0.34 | 0.739 | -15.55 | 11.14 | 0.003 |
| **Depression Symptoms** | -0.07 | 0.57 | -0.12 | 0.901 | -1.23 | 1.08 | 0.000 |
| **Twin Relatedness** | -0.33 | 0.36 | -0.94 | 0.355 | -1.05 | 0.39 | 0.024 |
| **AU Group** | -10.05 | 6.04 | -1.66 | 0.105 | -22.30 | 2.21 | 0.073 |

**miR-150-5p**

| Parameter | B | SE | t | *p* value | 95% CI | | Partial $\eta^2$ |
|---|---|---|---|---|---|---|---|
| | | | | | Lower Bound | Upper Bound | |
| **Sex** | 3.28 | 2.38 | 1.37 | 0.178 | -1.56 | 8.12 | 0.051 |
| **Age** | -0.06 | 1.12 | -0.05 | 0.958 | -2.33 | 2.21 | 0.000 |
| **Smoking Behaviors** | 2.62 | 3.24 | 0.81 | 0.423 | -3.95 | 9.19 | 0.018 |
| **Depression Symptoms** | 0.48 | 0.28 | 1.72 | 0.094 | -0.09 | 1.05 | 0.078 |
| **Twin Relatedness** | 0.02 | 0.17 | 0.12 | 0.906 | -0.33 | 0.38 | 0.000 |
| **AU Group** | -1.43 | 2.97 | -0.48 | 0.634 | -7.46 | 4.61 | 0.007 |

**miR-181a-5p**

| Parameter | B | SE | t | *p* value | 95% CI | | Partial $\eta^2$ |
|---|---|---|---|---|---|---|---|
| | | | | | Lower Bound | Upper Bound | |
| **Sex** | -0.35 | 0.21 | -1.65 | 0.107 | -0.79 | 0.08 | 0.075 |
| **Age** | 0.01 | 0.10 | 0.06 | 0.955 | -0.20 | 0.21 | 0.000 |
| **Smoking Behaviors** | -0.18 | 0.29 | -0.62 | 0.541 | -0.76 | 0.41 | 0.011 |
| **Depression Symptoms** | 0.01 | 0.02 | 0.26 | 0.798 | -0.04 | 0.06 | 0.002 |
| **Twin Relatedness** | -0.01 | 0.02 | -0.42 | 0.681 | -0.04 | 0.03 | 0.005 |
| **AU Group** | -0.06 | 0.27 | -0.24 | 0.815 | -0.62 | 0.49 | 0.002 |

**miR-199a-5p**

| Parameter | B | SE | t | *p* value | 95% CI | | Partial $\eta^2$ |
|---|---|---|---|---|---|---|---|
| | | | | | Lower Bound | Upper Bound | |
| **Sex** | 0.40 | 0.91 | 0.44 | 0.659 | -1.44 | 2.25 | 0.006 |
| **Age** | -0.21 | 0.43 | -0.49 | 0.628 | -1.07 | 0.66 | 0.007 |
| **Smoking Behaviors** | 0.30 | 1.23 | 0.24 | 0.808 | -2.20 | 2.80 | 0.002 |
| **Depression Symptoms** | -0.06 | 0.11 | -0.53 | 0.599 | -0.27 | 0.16 | 0.008 |
| **Twin Relatedness** | -0.01 | 0.07 | -0.17 | 0.869 | -0.15 | 0.12 | 0.001 |
| **AU Group** | 1.20 | 1.13 | 1.06 | 0.294 | -1.09 | 3.50 | 0.031 |

**miR-200c-3p**

| Parameter | B | SE | t | *p* value | 95% CI | | Partial $\eta^2$ |
|---|---|---|---|---|---|---|---|
| | | | | | Lower Bound | Upper Bound | |
| **Sex** | 0.13 | 0.11 | 1.21 | 0.235 | -0.09 | 0.35 | 0.040 |
| **Age** | -0.05 | 0.05 | -0.95 | 0.348 | -0.15 | 0.06 | 0.025 |
| **Smoking Behaviors** | 0.31 | 0.15 | 2.08 | 0.045 | 0.01 | 0.61 | 0.110 |
| **Depression Symptoms** | 0.02 | 0.01 | 1.60 | 0.118 | -0.01 | 0.05 | 0.068 |
| **Twin Relatedness** | 0.00 | 0.01 | -0.43 | 0.669 | -0.02 | 0.01 | 0.005 |
| **AU Group** | 0.25 | 0.14 | 1.81 | 0.079 | -0.03 | 0.52 | 0.085 |

**miR-30d-5p**

| Parameter | B | SE | t | *p* value | 95% CI | | Partial $\eta^2$ |
|---|---|---|---|---|---|---|---|
| | | | | | Lower Bound | Upper Bound | |
| **Sex** | 0.27 | 1.57 | 0.17 | 0.865 | -2.92 | 3.46 | 0.001 |
| **Age** | -0.16 | 0.74 | -0.22 | 0.826 | -1.66 | 1.33 | 0.001 |
| **Smoking Behaviors** | -0.25 | 2.13 | -0.12 | 0.908 | -4.58 | 4.08 | 0.000 |
| **Depression Symptoms** | -0.11 | 0.18 | -0.57 | 0.571 | -0.48 | 0.27 | 0.009 |
| **Twin Relatedness** | -0.01 | 0.12 | -0.11 | 0.912 | -0.25 | 0.22 | 0.000 |
| **AU Group** | 2.64 | 1.96 | 1.35 | 0.186 | -1.34 | 6.62 | 0.049 |

**miR-324-5p**

| Parameter | B | SE | t | *p* value | 95% CI | | Partial $\eta^2$ |
|---|---|---|---|---|---|---|---|
| | | | | | Lower Bound | Upper Bound | |
| **Sex** | 0.10 | 0.29 | 0.34 | 0.735 | -0.49 | 0.68 | 0.003 |
| **Age** | -0.03 | 0.14 | -0.20 | 0.843 | -0.30 | 0.25 | 0.001 |
| **Smoking Behaviors** | 0.01 | 0.39 | 0.03 | 0.974 | -0.78 | 0.81 | 0.000 |
| **Depression Symptoms** | 0.00 | 0.03 | -0.13 | 0.899 | -0.07 | 0.06 | 0.000 |
| **Twin Relatedness** | 0.01 | 0.02 | 0.25 | 0.803 | -0.04 | 0.05 | 0.002 |
| **AU Group** | 0.40 | 0.36 | 1.11 | 0.277 | -0.33 | 1.13 | 0.034 |

**miR-328-3p**

| Parameter | B | SE | t | *p* value | 95% CI | | Partial $\eta^2$ |
|---|---|---|---|---|---|---|---|
| | | | | | Lower Bound | Upper Bound | |
| **Sex** | 0.71 | 0.77 | 0.92 | 0.366 | -0.86 | 2.27 | 0.023 |
| **Age** | 0.09 | 0.36 | 0.24 | 0.814 | -0.65 | 0.82 | 0.002 |
| **Smoking Behaviors** | -0.35 | 1.05 | -0.34 | 0.738 | -2.48 | 1.77 | 0.003 |
| **Depression Symptoms** | -0.01 | 0.09 | -0.13 | 0.895 | -0.20 | 0.17 | 0.001 |
| **Twin Relatedness** | -0.04 | 0.06 | -0.78 | 0.439 | -0.16 | 0.07 | 0.017 |
| **AU Group** | -1.34 | 0.96 | -1.40 | 0.171 | -3.30 | 0.61 | 0.053 |

**miR-425-5p**

| Parameter | B | SE | t | *p* value | 95% CI |
|---|---|---|---|---|---|

| | | | | | 95% CI | | |
|---|---|---|---|---|---|---|---|
| | | | | | Lower Bound | Upper Bound | Partial $\eta^2$ |
| **Sex** | -0.24 | 0.24 | -1.02 | 0.316 | -0.72 | 0.24 | 0.029 |
| **Age** | -0.12 | 0.11 | -1.06 | 0.296 | -0.34 | 0.11 | 0.031 |
| **Smoking Behaviors** | -0.14 | 0.32 | -0.44 | 0.660 | -0.79 | 0.51 | 0.006 |
| **Depression Symptoms** | -0.01 | 0.03 | -0.50 | 0.620 | -0.07 | 0.04 | 0.007 |
| **Twin Relatedness** | -0.01 | 0.02 | -0.56 | 0.579 | -0.04 | 0.03 | 0.009 |
| **AU Group** | 0.04 | 0.29 | 0.15 | 0.883 | -0.55 | 0.64 | 0.001 |

**miR-501-3p**

| Parameter | B | SE | t | *p* value | 95% CI | | Partial $\eta^2$ |
|---|---|---|---|---|---|---|---|
| | | | | | Lower Bound | Upper Bound | |
| **Sex** | 0.01 | 0.01 | 2.13 | 0.041 | 0.00 | 0.03 | 0.118 |
| **Age** | 0.00 | 0.00 | 1.04 | 0.308 | 0.00 | 0.01 | 0.031 |
| **Smoking Behaviors** | 0.01 | 0.01 | 0.56 | 0.580 | -0.01 | 0.02 | 0.009 |
| **Depression Symptoms** | 0.00 | 0.00 | -0.64 | 0.529 | 0.00 | 0.00 | 0.012 |
| **Twin Relatedness** | 0.00 | 0.00 | 2.14 | 0.039 | 0.00 | 0.00 | 0.119 |
| **AU Group** | 0.00 | 0.01 | 0.29 | 0.775 | -0.02 | 0.02 | 0.002 |

**miR-574-3p**

| Parameter | B | SE | t | *p* value | 95% CI | | Partial $\eta^2$ |
|---|---|---|---|---|---|---|---|
| | | | | | Lower Bound | Upper Bound | |
| **Sex** | 0.16 | 0.11 | 1.45 | 0.155 | -0.06 | 0.39 | 0.057 |
| **Age** | -0.03 | 0.05 | -0.50 | 0.617 | -0.13 | 0.08 | 0.007 |
| **Smoking Behaviors** | 0.10 | 0.15 | 0.68 | 0.498 | -0.20 | 0.41 | 0.013 |
| **Depression Symptoms** | 0.01 | 0.01 | 0.53 | 0.601 | -0.02 | 0.03 | 0.008 |
| **Twin Relatedness** | 0.00 | 0.01 | -0.32 | 0.754 | -0.02 | 0.01 | 0.003 |
| **AU Group** | 0.05 | 0.14 | 0.36 | 0.719 | -0.23 | 0.33 | 0.004 |

**miR-874-3p**

| Parameter | B | SE | t | *p* value | 95% CI | | Partial $\eta^2$ |
|---|---|---|---|---|---|---|---|
| | | | | | Lower Bound | Upper Bound | |

| | | | | | | | |
|---|---|---|---|---|---|---|---|
| **Sex** | 0.06 | 0.10 | 0.62 | 0.539 | -0.14 | 0.26 | 0.011 |
| **Age** | -0.04 | 0.05 | -0.83 | 0.410 | -0.13 | 0.06 | 0.019 |
| **Smoking Behaviors** | 0.14 | 0.13 | 1.05 | 0.301 | -0.13 | 0.41 | 0.030 |
| **Depression Symptoms** | 0.00 | 0.01 | 0.25 | 0.807 | -0.02 | 0.03 | 0.002 |
| **Twin Relatedness** | 0.00 | 0.01 | -0.02 | 0.984 | -0.01 | 0.01 | 0.000 |
| **AU Group** | 0.12 | 0.12 | 0.95 | 0.347 | -0.13 | 0.37 | 0.025 |

**miR-877-5p**

| **Parameter** | **B** | **SE** | **t** | ***p* value** | **95% CI** | | **Partial $\eta^2$** |
|---|---|---|---|---|---|---|---|
| | | | | | **Lower Bound** | **Upper Bound** | |
| **Sex** | -0.09 | 0.12 | -0.76 | 0.451 | -0.34 | 0.16 | 0.016 |
| **Age** | -0.05 | 0.06 | -0.86 | 0.396 | -0.17 | 0.07 | 0.021 |
| **Smoking Behaviors** | 0.18 | 0.17 | 1.05 | 0.299 | -0.16 | 0.52 | 0.031 |
| **Depression Symptoms** | 0.00 | 0.01 | 0.29 | 0.773 | -0.03 | 0.03 | 0.002 |
| **Twin Relatedness** | 0.00 | 0.01 | -0.35 | 0.728 | -0.02 | 0.02 | 0.004 |
| **AU Group** | 0.24 | 0.15 | 1.58 | 0.122 | -0.07 | 0.55 | 0.067 |

**Figure S1. Potential internal controls normalized by miR-133a-3p**

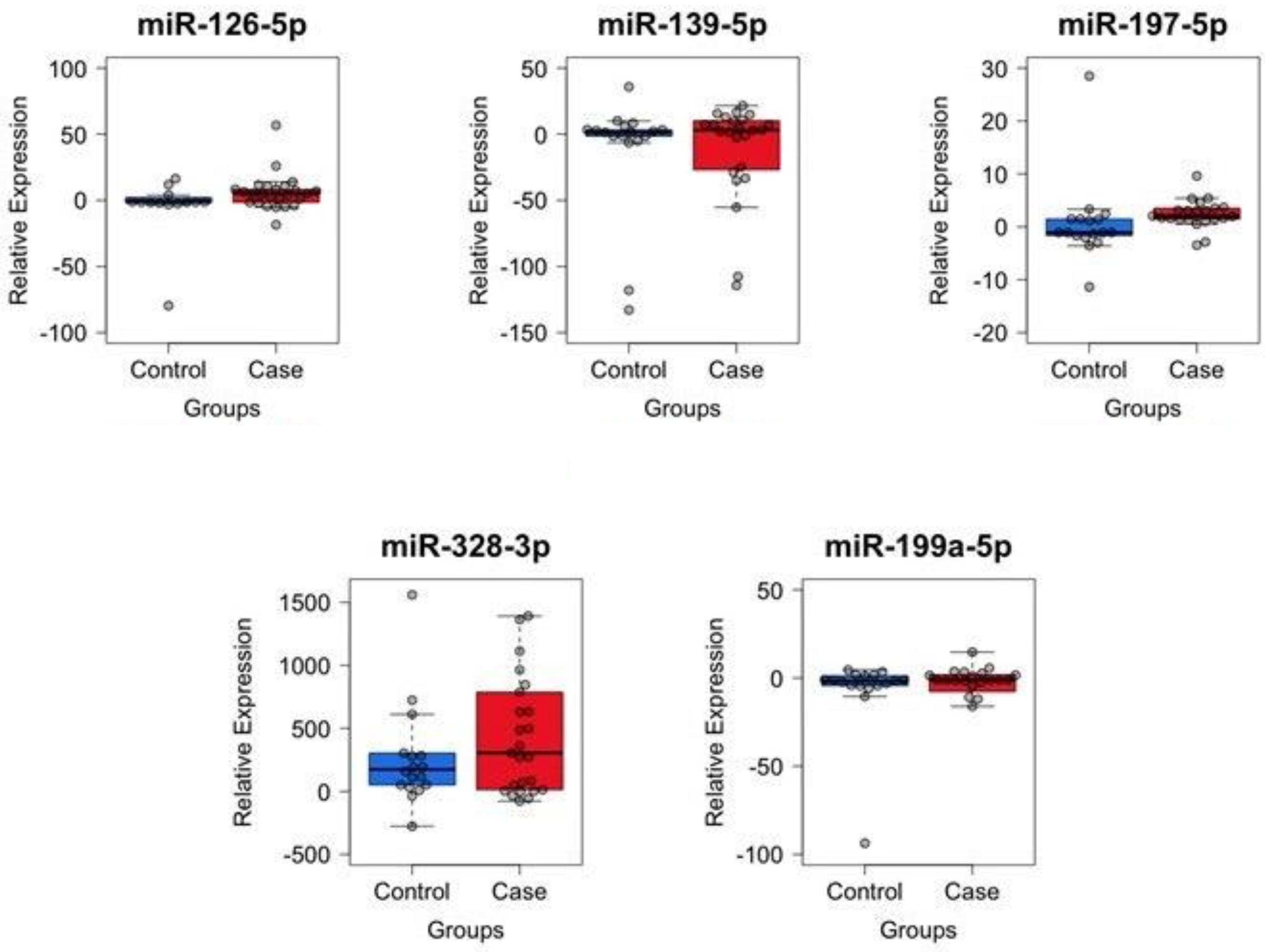

**Figure S2. qPCR analysis of individual samples from alcohol consumption (Case) and control groups.**

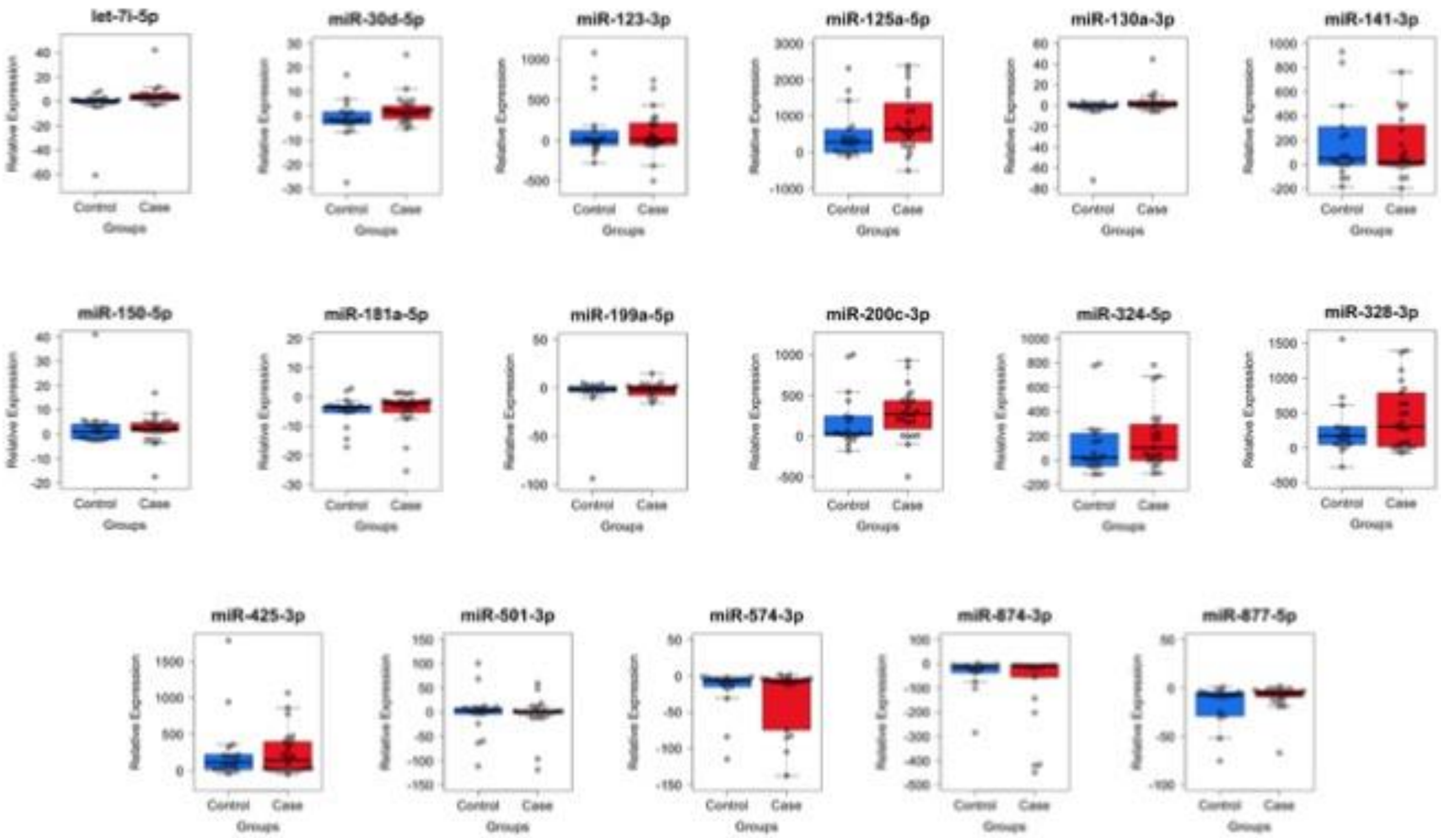
